# Computational prediction of MHC anchor locations guide neoantigen identification and prioritization

**DOI:** 10.1101/2020.12.08.416271

**Authors:** Huiming Xia, Joshua McMichael, Michelle Becker-Hapak, Onyinyechi C. Onyeador, Rico Buchli, Ethan McClain, Patrick Pence, Suangson Supabphol, Megan M. Richters, Anamika Basu, Cody A. Ramirez, Cristina Puig-Saus, Kelsy C. Cotto, Sharon L. Freshour, Jasreet Hundal, Susanna Kiwala, S. Peter Goedegebuure, Tanner M. Johanns, Gavin P. Dunn, Antoni Ribas, Christopher A. Miller, William E. Gillanders, Todd A. Fehniger, Obi L. Griffith, Malachi Griffith

## Abstract

Neoantigens are novel peptide sequences resulting from sources such as somatic mutations in tumors. Upon loading onto major histocompatibility complex (MHC) molecules, they can trigger recognition by T cells. Accurate neoantigen identification is thus critical for both designing cancer vaccines and predicting response to immunotherapies. Neoantigen identification and prioritization relies on correctly predicting whether the presenting peptide sequence can successfully induce an immune response. As the majority of somatic mutations are single nucleotide variants, changes between wildtype and mutated peptides are typically subtle and require cautious interpretation. A potentially underappreciated variable in neoantigen-prediction pipelines is the mutation position within the peptide relative to its anchor positions for the patient’s specific MHC molecules. While a subset of peptide positions are presented to the T-cell receptor for recognition, others are responsible for anchoring to the MHC, making these positional considerations critical for predicting T-cell responses. We computationally predicted high probability anchor positions for different peptide lengths for 328 common HLA alleles and identified unique anchoring patterns among them. Analysis of 923 tumor samples shows that 6-38% of neoantigen candidates are potentially misclassified and can be rescued using allelespecific knowledge of anchor positions. A subset of anchor results were orthogonally validated using protein crystallography structures. Representative anchor trends were experimentally validated using peptide-MHC stability assays and competition binding assays. By incorporating our anchor prediction results into neoantigen prediction pipelines, we hope to formalize, streamline and improve the identification process for relevant clinical studies.

**One Sentence Summary:** Neoantigen prediction accuracy is significantly influenced by the mutation position within the neoantigen and its relative position to the patient’s allele-specific MHC anchor locations.

## Introduction

Neoantigens arise from short peptide sequences specifically found in tumor cell populations resulting from sources such as somatic mutations, RNA editing*(1)*, alternative splicing*(2)*, etc. They can be loaded onto major histocompatibility complex (MHC) class I or II molecules to allow recognition by cytotoxic or helper T cells. Upon recognition, T cells are then able to signal cell death for an anti-tumor response. Multiple studies have shown the potential of neoantigen based immunotherapy treatments for cancer*(3–5)* and numerous clinical trials are underway. Accurate neoantigen prediction and prioritization is critical to understanding tumor immunology, response to checkpoint blockade therapy, and for the design of personalized vaccines*(6)* and T cell therapies. Several bioinformatic tools and pipelines have been developed to facilitate neoantigen identification*(7–10)*.

The effectiveness of a neoantigen-based vaccine relies in part on whether the neoantigen sequences presented to T cells have previously been exposed to the immune system and would be subject to central tolerance (where immune response to antigens is limited as a result of clonal deletion of autoreactive B cells and T cells). While a variety of mutation types are being explored as neoantigen sources*(11–15)*, the vast majority of somatic mutations currently identified as sources of neoantigens are single nucleotide variants (SNVs)*(4, 16)*, though emerging research suggests that we are underestimating the potential for neoantigen generation from more complex forms of variation *(17–20)*. Amino acid sequence changes between the wildtype (WT) and mutant (MT) peptides are subtle (often a single amino acid substitution) and mutant peptides remain similar to native sequences of the host. Additionally, only a subset of positions on the loaded peptide sequence are primarily presented to the T-cell receptor for recognition, and another subset of positions are primarily responsible for anchoring to the MHC, making these positional considerations critical for predicting T-cell responses (**Fig. 1**). Thus, subtle amino acid changes must be interpreted cautiously. Multiple factors should be considered when prioritizing neoantigens, including mutation location, anchor position, predicted MT and WT binding affinities, and WT/MT fold change, also known as agretopicity*(21)*. Examples of the four distinct possible scenarios for a predicted strong MHC binding peptide involving these factors are illustrated in **Fig. 1**. There are other possible scenarios where the MT is a poor binder, however those are not listed as they would not pertain to our goal of neoantigen identification. The first scenario shows the cases where the WT is a poor binder and the MT peptide, a strong binder, contains a mutation at an anchor location. Here, the mutation results in a tighter binding of the MHC and allows for better presentation and potential for recognition by the TCR. As the WT does not bind (or is a poor binder), this neoantigen remains a good candidate since the sequence presented to the TCR is novel. The second and third scenarios both have strong binding WT and MT peptides. In the second scenario, the mutation of the peptide is located at a non-anchor location, creating a difference in the sequence participating in TCR recognition compared to the WT sequence. In this case, although the WT is a strong binder, the neoantigen remains a good candidate that should not be subject to central tolerance. However, as shown in the third scenario, there are neoantigen candidates where the mutation is located at the anchor position and both peptides are strong binders. Although anchor positions can themselves influence TCR recognition*(22)*, a mutation at a strong anchor location generally implies that both WT and MT peptides will present the same residues for TCR recognition. As the WT peptide is a strong binder, the MT neoantigen, while also a strong binder, will likely be subject to central tolerance and should not be considered for prioritization. The last scenario is similar to the first scenario where the WT is a poor binder. However, in this case, the mutation is located at a non-anchor position, likely resulting in a different set of residues presented to the TCR and thus making the neoantigen a good candidate. Recent studies on neoantigens for both mouse and human models have confirmed the importance of anchor location when predicting the overall immunogenicity of a given peptide*(23, 24)*. However, it is important to note that these scenarios are not absolute. While a position that acts as a strong anchor is less likely to be TCR facing (and vice versa), certain positions of the peptide may interact to varying degrees with both the MHC and TCR.

**Fig. 1.**
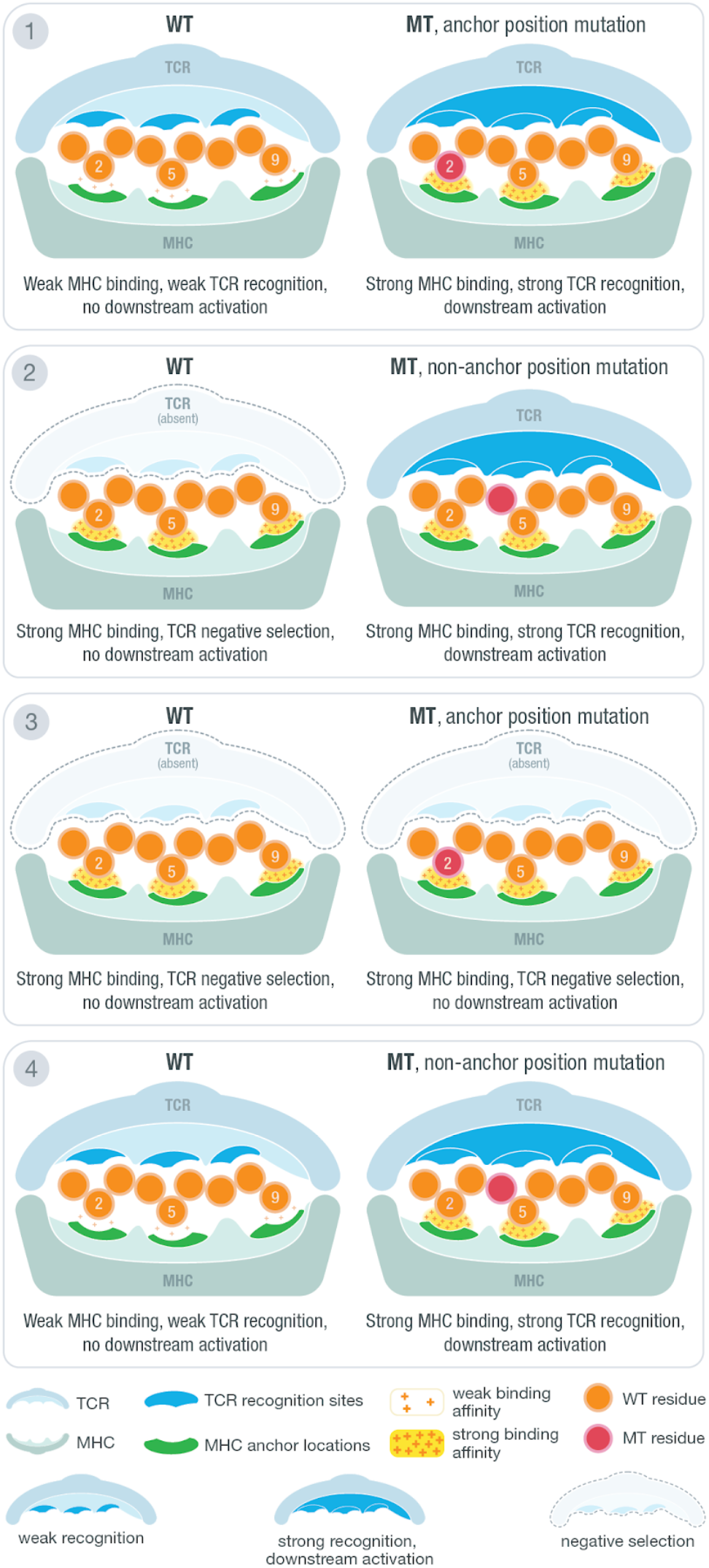
Anchor and mutation position scenarios at the MHC-peptide-TCR interface. Illustration of MHC-peptide-TCR interface using an example structure with anchors at position 2, 5 and 9. At the contact interface between the peptide-loaded MHC and the recognizing T-cell receptor, certain positions are responsible for anchoring the peptide to the MHC molecule and/or potentially being recognized by the TCR. The position of tumor specific (“mutant”) amino acids relative to anchor positions and predicted binding affinity of mutant and wild type peptides produce four distinct scenarios for interpreting candidate neoantigens. Example TCR recognition sites are shown in blue while MHC anchor locations are shown in green. The peptide residues are shown in orange while the mutant residue is marked with red. A yellow force field with varying density is used to illustrate binding strength between peptide and the MHC molecule. Three different cases of TCR recognition level are depicted including: self-recognizing TCR absent due to negative selection, weak-recognizing TCR due to weak MHC binding of presented peptide, and strong-recognition of TCR triggering downstream activation of cytotoxic T-cells.

The mutation’s position within the peptide relative to its anchor positions for the patient’s human leukocyte antigen (HLA) alleles, is currently overlooked by neoantigen prediction pipelines. However, failing to account for these positional considerations may result in susceptibility to central tolerance and potentially induce auto-immunity. Many recently published neoantigen studies have used simple filtering strategies with either only binding affinity filters*(5, 25)* (e.g. MT peptide IC50 < 500 nM) or with an additional agretopicity filter*(26–28)*, all without specifying whether they account for anchor and mutation locations during their selection process. Researchers have previously discussed how anchor locations can affect our interpretation of other factors considered in neoantigen prioritization (e.g. MT, WT binding affinities)*(29)*. However, a systematic method for determining anchor locations for the wide range of HLA alleles present in the population and application of these to evaluate MT/WT peptide pairs arising in tumors has not been reported. As a result, many neoantigen studies have either failed to adequately consider this crucial factor or have used simplified and arbitrary assumptions to guide their neoantigen identification process.

Here, we provide a computational workflow for predicting anchor locations for a wide range of HLA alleles using a seed dataset generated from a collection of patient samples from local tumor sequences studies*(30)* combined with samples from The Cancer Genome Atlas (TCGA). Analysis of results showed clusters of different anchor trends among the HLA alleles analyzed and a subset of these HLA anchor results were orthogonally validated using protein crystallography structures. Representative anchor trends were experimentally validated using cell-based stability assays and IC50 binding assays. Using additional TCGA samples, we further evaluated how prioritization results may change when provided with additional anchor information, emulating a subset of steps in the neoantigen selection process by an immunotherapy tumor board, tasked with prioritizing vaccine candidates. By sharing our results for incorporation into neoantigen prediction pipelines, we hope to improve neoantigen prioritization for relevant experimental and clinical studies.

## Results

### Computational and quantitative prediction of HLA-specific anchor positions

In order to predict anchor locations for a wide range of HLA alleles, we assembled a seed HLA-peptide dataset of strong binding peptides with a median predicted IC50 of less than 500 nM across (up to) 8 MHC class I algorithms (NetMHC*(31)*, NetMHCpan*(32)*, MHCnuggets*(33)*, MHCflurry*(34)*, SMM*(35)*, Pickpocket*(36)*, SMMPMBEC*(37)*, NetMHCcons*(38)*). These peptides were obtained from TCGA and supplemented with additional patient datasets from our own neoantigen study cohorts including lymphoma, glioblastoma*(30)*, breast cancer, and melanoma (**Methods; Data file S1**). A total of 609,807 peptides were identified with the majority being 9-mers and 10-mers (**Fig. S1**). In total, these peptides corresponded to 328 HLA alleles across 1,443 tumor samples.

For each individual HLA allele of which data was obtained, peptides were sorted and separated by their respective lengths, ranging from 8 to 11 (**Fig. 2**). These peptides were then mutated in silico at all possible positions to all possible amino acids. Predicted binding affinities for each individual peptide were then obtained using the same set of algorithms as previously described. These binding affinities were compared to the median binding affinity of the original strong binding peptide sequence. This comparison enables us to evaluate how mutations occurring at each individual position change the predicted binding interaction between the strong binding peptide and the MHC molecule. A significant change observed at a particular location indicates a higher probability of the amino acid at the position acting as an anchor. On the other hand, little to no change in binding affinities when a position is mutated would indicate a lower probability of the position acting as an anchor. An overall score per position was obtained by summing across all peptides analyzed for an individual HLA allele (**Fig. 2; Fig. S2; Methods)**.

**Fig. 2.**
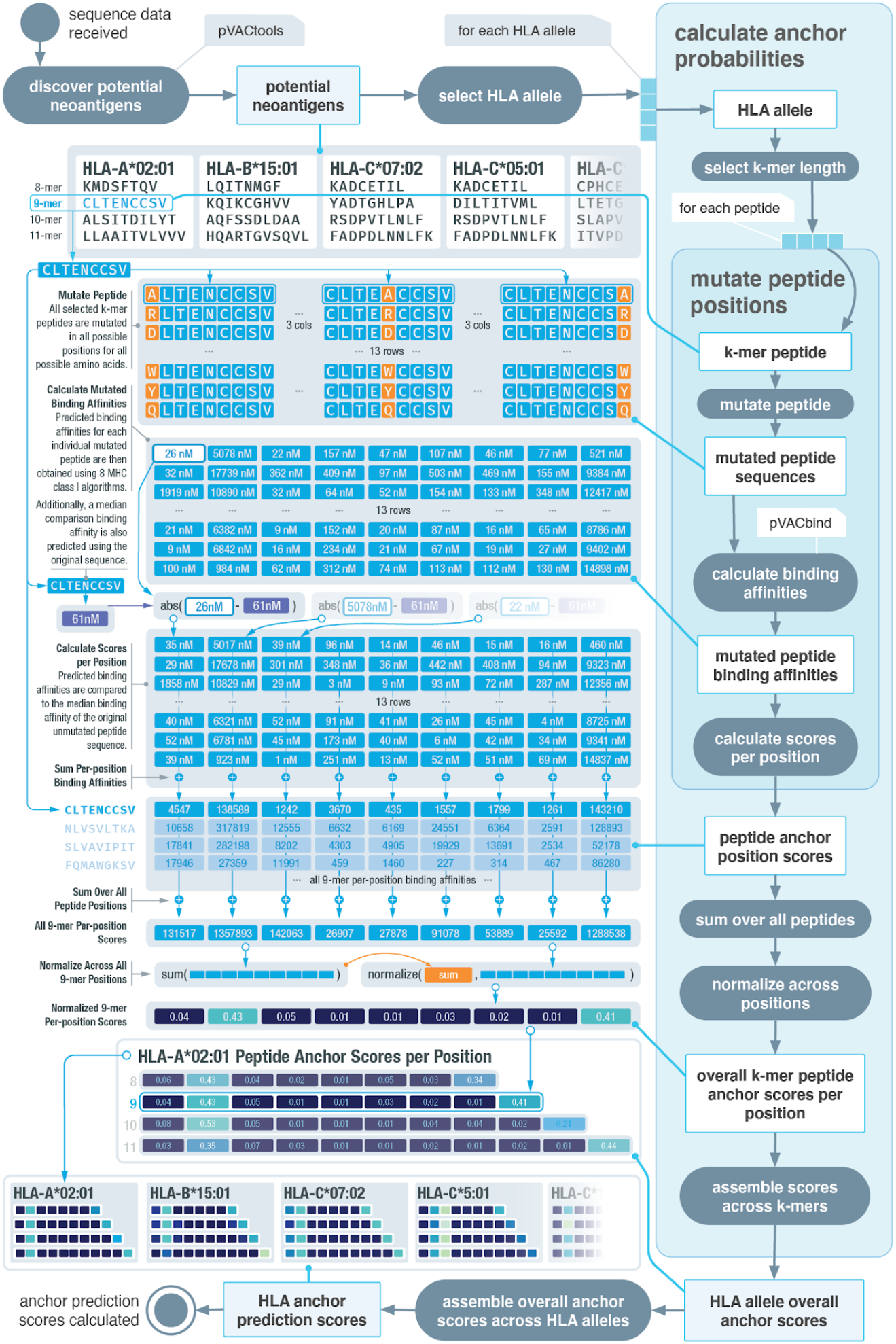
Overview of computational workflow for anchor position prediction. Schematic of our computational workflow to simulate the effect of mutation position on MHC binding affinities of peptides for individual HLA alleles. HLA and peptide pairings are selected from a reference dataset of putative strong binders. All possible amino acid changes are applied to all possible positions and the impact on binding affinity is assessed. An overall peptide anchor score is calculated for each position for all HLA-peptide length combinations. Higher scores indicate greater likelihood that a particular position in the peptide acts as an anchor residue for a given HLA.

### Prediction results show distinct patterns of HLA anchor locations

We generated anchor prediction scores for the 328 HLA alleles with strong binding peptides in our seed dataset (**Data file S2**). These HLA alleles include 95 HLA-A alleles (representing 99.2% of the HLA-A alleles observed in the population according to the Allele Frequency Net Database), 175 HLA-B alleles (representing 97.9% of HLA-B alleles) and 58 HLA-C alleles (representing 98.5% of the HLA-C alleles)*(39)*(**Methods**). Results were separated based on peptide lengths (8–11) and the anchor prediction scores across all HLA alleles were visualized using hierarchical clustering with average linkage (**Fig. 3; Fig. S3**). We observed different anchor patterns across HLA alleles, varying in both the number of anchor positions as well as the location. These anchor position patterns could be roughly clustered into 6 distinct groups. Additionally, while we utilized peptides generated from cancer patient data, we wanted to see if changing the source of the peptides would affect our predicted anchor patterns (**Fig. S4; Methods**). After averaging calculations over 1,000 peptides from each source, we observed high convergence of the anchor pattern for HLA-A*02:01 across all four different sources (random, reference proteome, viral proteome, and cancer mutations). This demonstrated the possibility of expanding results to a wider span of HLA alleles in the future by utilizing different sources of peptides.

**Fig. 3.**
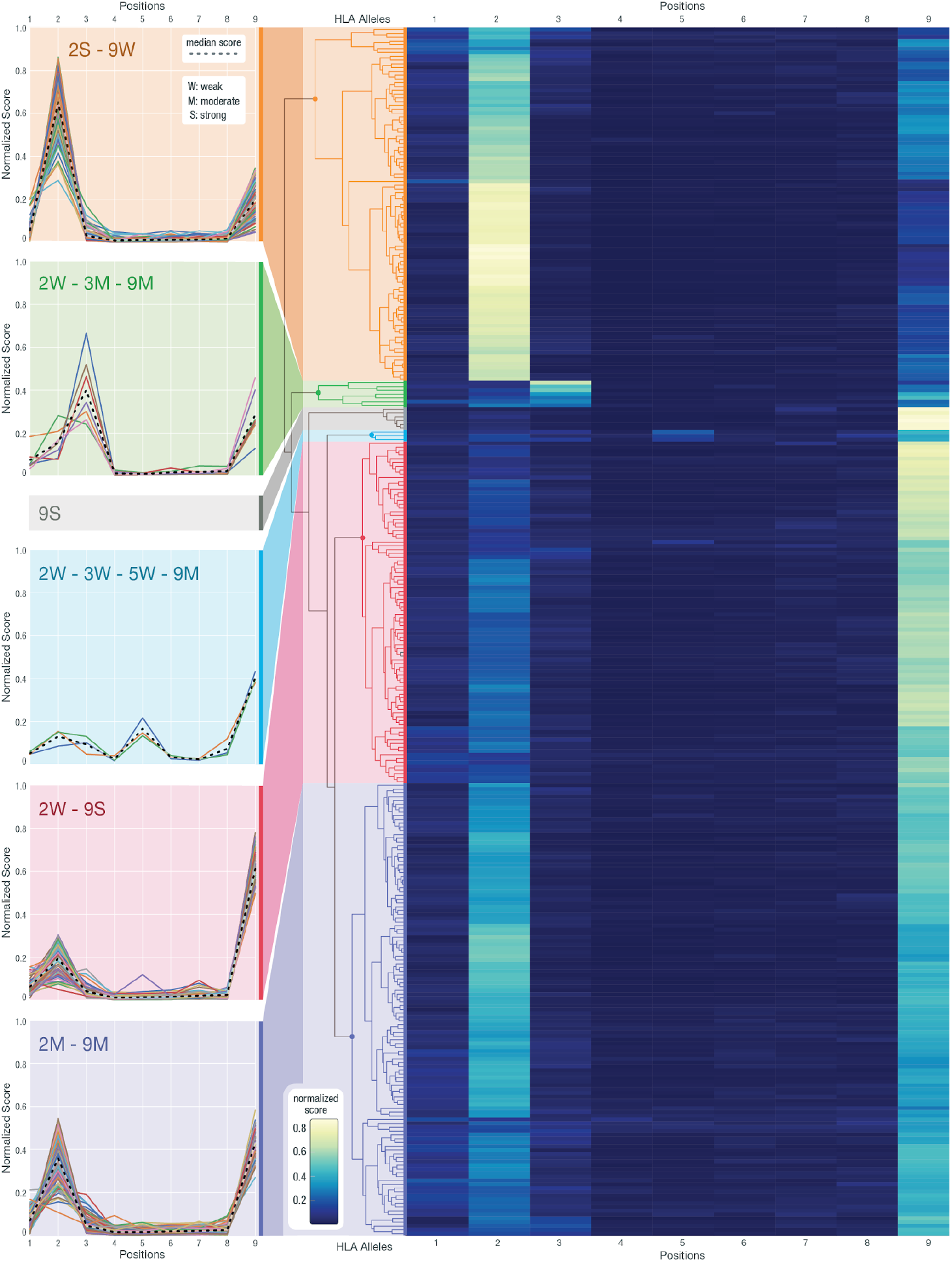
Hierarchical clustering of anchor prediction scores across all 9-mer peptides. Anchor prediction scores clustered using hierarchical clustering with average linkage for all 318 HLA alleles for which 9-mer peptide data were collected (out of the 328 HLA alleles, 10 did not have corresponding 9-mer data). For the heatmap, the x-axis represents the 9 peptide positions and the y-axis represents 318 HLA alleles. Example HLA clusters have been highlighted with various color bands and the score trends for individual HLA alleles are plotted. In the cluster line plots on the left, the x-axis shows the peptide positions while the y-axis corresponds to the anchor score, normalized across all peptide positions. Different annotations have been given to help summarize the trends observed in individual clusters, where numbers represent positions and letters represent its strength as a potential anchor in comparison to other anchors (S: strong, M: moderate, W: weak). The median scores for each cluster are presented with a dashed line.

Previously, anchor locations have generally been assumed to be at the second and terminal position of the peptide with equal weighting (with the notable exception of HLA-B*08:01)*(40)*. Our 9-mer clustering results confirm that the majority of HLA alleles predicted show positions 2 and 9 as likely anchor locations (95% fall into the 2S-9W, 2W-9S, 2M-9M clusters combined). However, three distinct cluster groups can be further identified within the larger group. The 2S-9W cluster represents HLA alleles with a strong anchor predicted at position 2 and a weak anchor predicted at position 9 (2S-9W; **Fig. 3**). The 2W-9S cluster shows those with a strong anchor predicted at position 9 and a weak anchor predicted at position 2 (2W-9S; **Fig. 3**). Additionally, we observe a smaller cluster of HLA alleles with moderate anchor predictions for both positions (2M-9M; **Fig. 3**) and another cluster with strong anchor predictions for only position 9 (9S; **Fig. 3**). We also discovered other patterns differing from the previous anchor assumptions of 2 and 9. In particular, we observed a clustered group of HLA-C alleles that have a moderate anchor at 3 and 9 accompanied by a weaker signal at 2 (2W-3M-9M; **Fig. 3**). A smaller group of HLA-B alleles also show an additional anchor at position 5 (2W-3W-5W-9M; **Fig. 3**). Our results indicate that a conventional anchor assumption putting equal weights on positions 2 and 9 does not capture the significant heterogeneity in anchor usage between different HLA alleles. These anchor considerations can affect neoantigen prioritization decisions and we hypothesized that HLA allele-specific anchor predictions would allow ranking of neoantigens with greater accuracy.

To analyze the possibility of our clusters arising due to bias from prediction algorithms and their training datasets, we examined the training dataset used by NetMHCpan4.0 for the 328 HLA alleles that we have included (**Data file S3**). Additionally, for alleles with limited data (hence predicted through a pan allele strategy), the ability to effectively utilize information from other alleles (for those lacking training data) also depends on how similar the allele of interest is to other alleles that do have robust amounts of data. Thus we also looked at the neighboring HLA alleles that are used in cases where training data was not available and the distance (calculated by NetMHCpan4.0) between each HLA allele and its nearest neighbor. Overall, the majority of HLA alleles (99.7%) had binding predictions from at least 4 different algorithms with 54 HLA alleles being supported by all 8 (**Fig. S5a**). The amount of training data for each HLA allele also correlated with how consistent binding predictions were across algorithms (**Fig. S5b**). This analysis confirms our expectation that more training data leads to more consistent predictions across algorithms. For the smaller clusters such as 2W-3W-5W-9M (blue), 2W-3W-9M (green) and 9S (gray), we can see that in general they fall along the same level as other clusters when comparing their nearest neighbor distances and how much variation is seen across prediction algorithms (**Fig. S5c,d**). Additionally, 2 out of the 3 alleles in the 2W-3W-5W-9M cluster, 4 out of 7 alleles in the 2W-3W-9M cluster and 6 out of 6 alleles in the 9S cluster all have some amount of training data available (**Fig. S5e, Data file S3**). When looking at network graphs showing HLA alleles and their neighbors, those with 2W-3W-5W-9M (blue) and 2W-3W-9M (green) patterns are scattered among different clusters with some alleles even acting as the center node (indicating non-zero amount of training data and thus no need to estimate binding based on similar alleles) (**Fig. S5f,g**). Thus, we concluded that the less common anchor clusters do not correlate with lack of training data nor similarity to a limited set of nearest neighbors.

Hence, we excluded the possibility of clusters arising due to bias in algorithms and their training data and believe that allele-specific differences are the true underlying reason for our observations.

### Protein structural analysis confirms predicted anchor results

To validate our anchor predictions, we collected X-ray crystallography structures for MHC molecules with bound peptides (**Data file S4**). The 166 protein structures collected corresponded to 33 HLA alleles with the majority of them containing 9-mer peptides (8-mers: 6, 9-mers: 110, 10-mers: 39, 11-mers: 11). These structures were analyzed using two methods: 1) measuring the physical distance between the peptide and the MHC binding groove and 2) calculating the solvent-accessible surface area (SASA) of the peptide residues (**Fig. 4a,b**; **Methods**). These methods were selected to validate predicted anchor positions based on the assumptions that if a certain peptide position is designated as an anchor then it is 1) more likely to be closer to the HLA molecule and 2) more secluded from solvent surrounding the peptide-MHC complex compared to non-anchor positions. This is because non-anchor peptide residues should be accessible to the TCR for recognition, thus in the peptide-MHC structures collected where a TCR is not present, peptide surface area available to the surrounding solvent roughly mimics the area that would be accessible by the TCR. Thus we expect an inverse correlation between our anchor prediction scores and the distance/SASA metrics. HLA-A*02:01, shown as an example, was found to have the greatest number of qualifying structures, with 47 of them containing a 9-mer peptide (**Fig. 4c**). These x-ray crystallography structures each capture a snapshot of a dynamic protein structure in constant movement. By overlaying the distance and SASA scores across all 47 complexes respectively, we observe that positions 2 and 9 are the ones most consistently close to the HLA molecule while also being secluded from the solvent. This observation corresponds well with our prediction of strong anchors at both positions 2 and 9 for HLA-A*02:01 and a 9-mer peptide. We also performed a targeted analysis on HLA-B*08:01 (2W-3W-5W-9M; blue cluster) with the limited data available and observed that positions 6 and 7 consistently bulged out whereas other positions tend to be closer to the HLA molecule while also being secluded from solvent (**Fig. S6**).

**Fig. 4.**
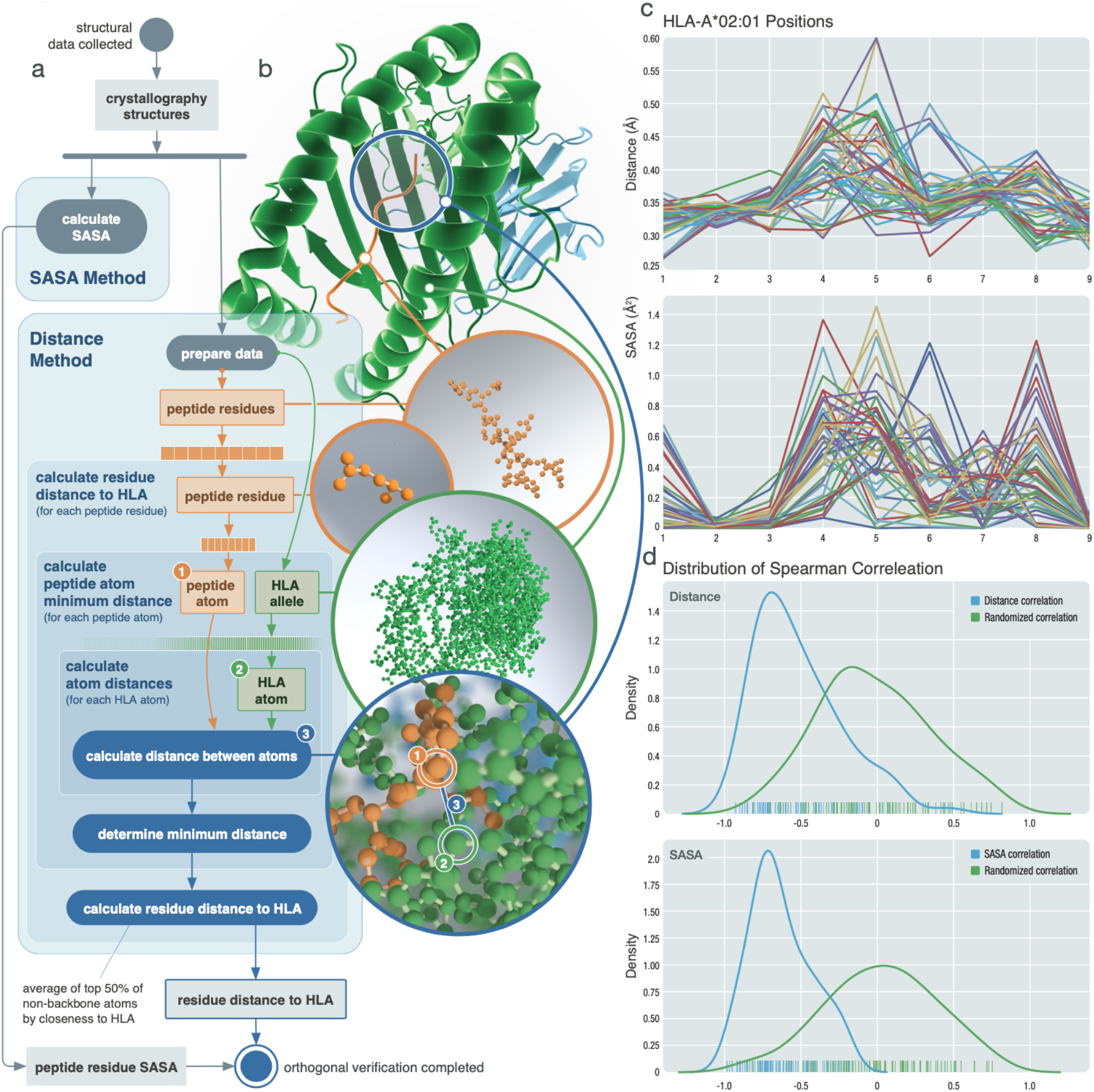
Orthogonal validation using protein crystallography structures. Orthogonal validation of predicted anchor scores utilizing X-ray crystallography structures. **a**, Schematic of analysis workflow for each HLA-peptide structure collected. For the distance metric, backbone atoms were excluded with the exception of glycine. **b**, Structural example of HLA-B*08:01 bound to peptide FLRGRAYGL (PDB ID: 3X13). **c**, Example results of 47 structures collected for HLA-A*02:01 with 9-mer peptides. Top panel corresponds to distance measurements for each position while the bottom panel corresponds to SASA measurements. **d**, Distribution of Spearman correlations calculated between distance and prediction scores (top) and SASA and prediction scores (bottom). Blue line represents each respective correlation distribution while the green line shows the distribution of Spearman correlation values obtained from randomly shuffled peptide positions.

To evaluate how the distance and SASA metric correlates with our prediction results across different HLA alleles, we calculated Spearman correlations between our prediction scores and distance/SASA results for each peptide position across 87 PDB structures. These structures were determined by randomly selecting at most 5 structures per HLA-length combination. The distribution of these correlations was compared to that of a randomized dataset where positions of the peptide were randomly shuffled (**Fig. 4d; Data file S5; Methods**). Two sample t-tests showed the distributions were significantly different from the randomized dataset with statistical values of −9.9795 (p value = 1.3757e-18) and −14.7322 (p value = 8.7472e-30) for distance and SASA metrics respectively. Results show that 91.95% of our prediction scores are inversely correlated with the distance metric and 100% of them are inversely correlated with the SASA scores. Furthermore, we analyzed 61 protein structures that contained both the peptide-MHC complex and an additional binding TCR molecule. The distance between the TCR and the peptide showed high correlation with our prediction scores (**Fig. S7**). Two-sample t-tests showed significant differences between the randomized dataset and both the HLA-peptide distance (p = 8.8023e-08) and the TCR-peptide distance (p=2.4915e-13). These results together strongly suggest that our anchor prediction workflow is accurately predicting allele specific anchor sites.

### Experimental validation shows similar anchor trends as our prediction across distinct HLA alleles

In order to further validate the predicted anchor patterns we observed, we selected a range of HLA alleles representing these patterns for experimental validation. We used two different experimental validation methods across our peptide-HLA combinations: in vitro IC50 binding assays and cell-based stabilization assays (**Methods)**. Validations were performed on a total of 136 peptide-HLA combinations across 8 different HLA alleles selected to represent varying anchor patterns. For each HLA allele selected, we first validated peptides that were predicted to be strong binders. These peptides were originally MT peptides from either clinical datasets or TCGA samples paired with matching HLA alleles from the patient. For a validated strong binding peptide-HLA combination, we then synthesized peptides mutated in multiple ways at different positions, which each had a varying predicted probability of acting as an anchor. These mutated peptides were then evaluated experimentally for MHC binding and/or stability and compared to the original strong-binding peptide (**Data file S6**). For example, we performed an in-depth analysis for HLA-B*07:02 with a strong binding 9-mer peptide RPDVKHSKM. Overall for 9-mer peptides, HLA-B*07:02 was predicted to have a high probability anchor at position 2 and a lower probability anchor at position 9 (2S-9W) (**Fig. 5a**). Following our computational prediction workflow, we also calculated the specific anchor trend for peptide RPDVKHSKM, which agrees with the overall 9-mer trend with positions 2, 9 and 1 as anchors in descending probability (**Fig. 5b**). We mutated positions 1, 2, 5 and 9 to four different amino acids, each with varying predicted influence on the binding affinity, and measured their stability with regards to HLA-B*07:02 (**Methods; Fig. 5c**). For non-anchor position P5, all mutated peptides remained strong binders. For the two weak anchor positions P1 and P9, certain amino acid changes disturbed the binding while others had little influence. For the strong anchor position P2, all amino acid changes led to a non-binder. We further performed IC50 binding assays and observed a similar trend to that of the stability experiments (**Fig. 5d**). Similar mutation experiments on HLA-B*08:01, HLA-A*68:01 and HLA-A*23:01 were conducted (**Fig. S8 and S9**). These additional experiments confirmed the varying strengths of individual positions acting as anchors for MHC alleles comprising anchor patterns 2W-3W-5W-9M, 2W-9S and 9S.

**Fig. 5.**
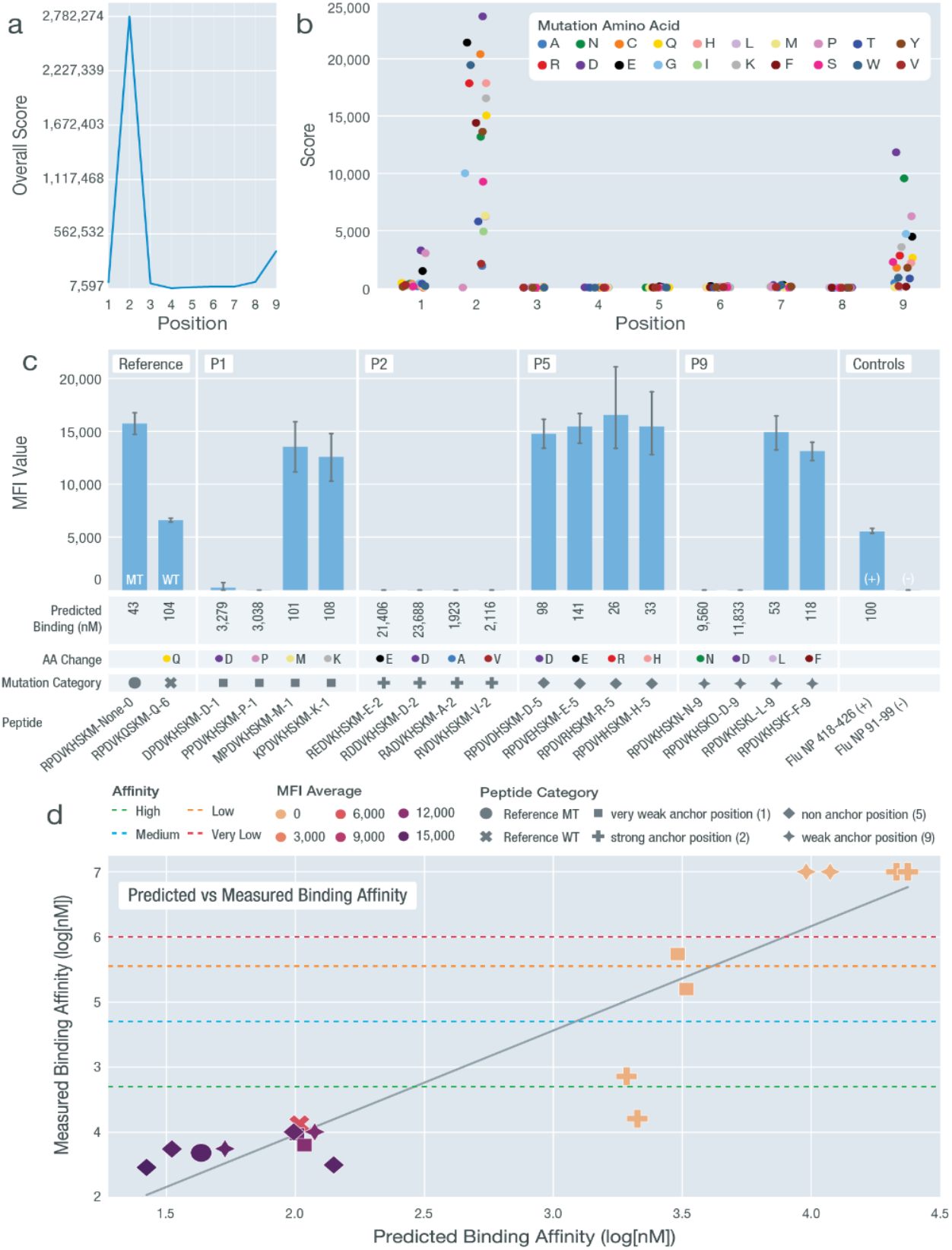
Experimental Validation of anchor pattern for HLA-B*07:02. Validation results for HLA-B*07:02 with a predicted 2S-9W anchor pattern using peptide sequence RPDVKHSKM. **a**, Overall anchor prediction scores for each position of 9-mer peptides when binding to HLA-B*07:02. Higher scores indicate a higher probability of acting as an anchor. **b**, Binding affinity changes (y-axis) plotted for each position of the specific 9-mer peptide RPDVKHSKM. Each peptide position was mutated to 19 other amino acids to evaluate influence on binding affinity. **c**, MFI values measured from cell stabilization assays are plotted for both unmutated and mutated peptides. Peptides are marked by their mutation positions (P1, P2, P5 and P9), predicted binding affinity values, amino acid changes (color coordinated with figure 5b), and mutation category (shape coordinated with figure 5d). **d**, Predicted binding affinity scores (log10[nM]) plotted against measured binding affinity values (log10[nM]) from IC50 binding assays. Binding categories based on measured binding affinity values are marked using horizontal lines. MFI values are overlaid using heatmap coloring for all data points. Peptide categories are marked using different shapes (coordinated with figure 5c). All nonbinders were plotted with a measured binding affinity value of 7.

Overall, across 136 peptide-HLA combinations, experimental validation results were consistent with the average predicted binding affinities calculated from 8 prediction algorithms. For peptides validated using IC50 binding assays, reasonable correlations were observed between the predicted and measured IC50 values, with respect to individual HLA alleles (**Fig. 6a**). When comparing the measured binding category (determined by the measured IC50) with predicted binding affinities, we saw a clear trend with high affinity binders having the lowest predicted IC50 values (**Fig. 6b**). A few outliers were observed where peptides were predicted as strong binders but when validating, these peptides were categorized as non-binding. This may be explained by difficulties in manufacturing and/or solubilizing these peptides, preventing detection in our validation experiments. As expected, an inverse correlation was observed between the MFI average values and the predicted binding affinities (**Fig. 6c**). Similarly, peptides with measured high affinities showed the highest MFI values. Peptides with measured binding affinities among the “medium”, “low”, “very low” and “no binder” categories were similar in their range of MFI values, indicating a difference in the sensitivity between the IC50 binding assays and cell stabilization experiments (**Fig. 6d; Methods**).

**Fig. 6.**
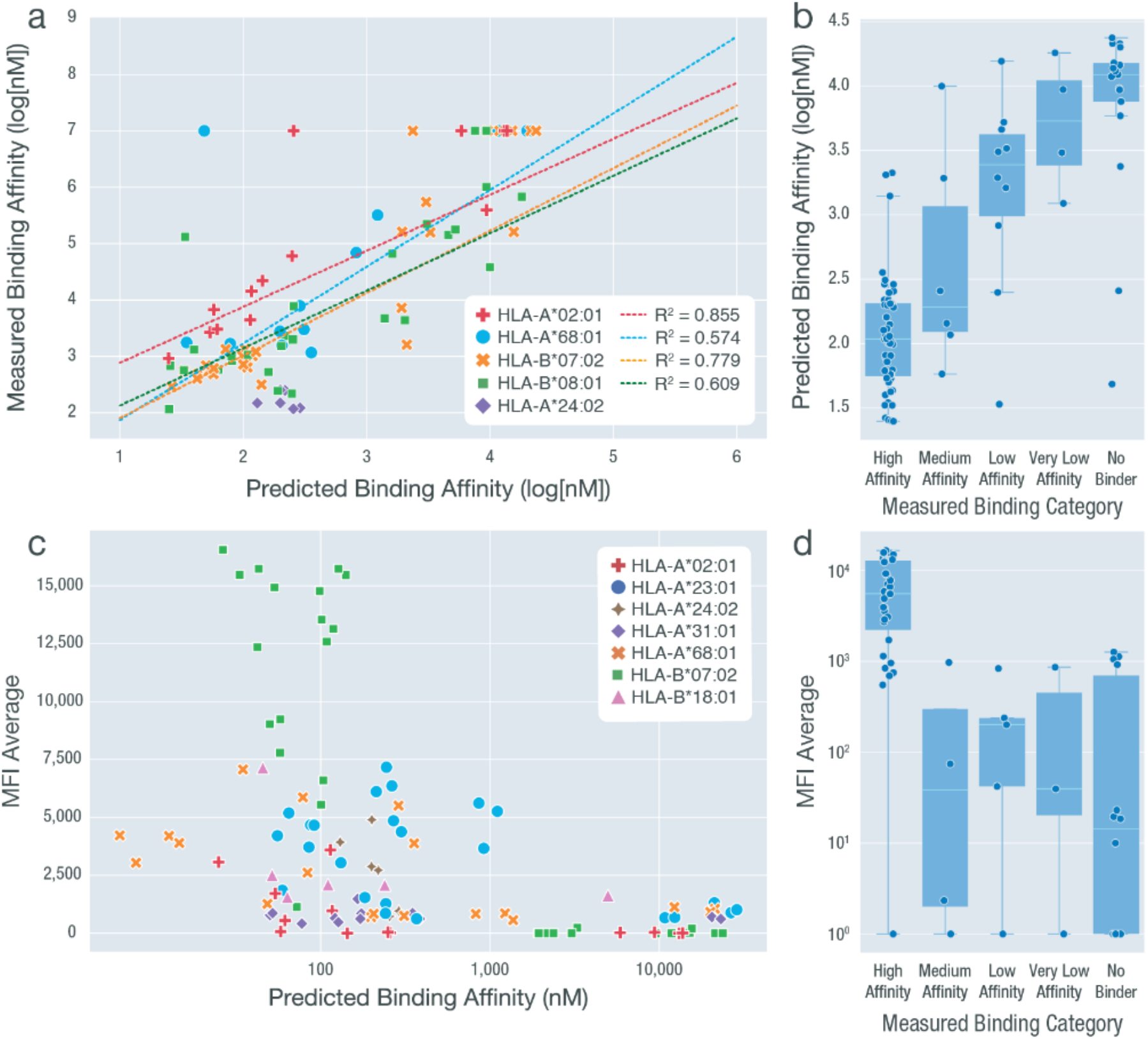
Summary of experimental validation results across all HLA allele-peptide combinations. We performed two types of experimental validations including cell-based stabilization assays (MFI values) and IC_50_ binding assays (log[nM]) for a total of 136 peptide-HLA combinations. The results from these assays are compared between each other and against the predicted binding affinity (averaged across 8 different algorithms). Measured binding categories are determined based on the measured IC_50_ values. **a**, Predicted binding affinity values plotted against the measured binding affinity (log[nM]). HLA-A*68:01, HLA-B*07:02, HLA-B*08:01 and HLA-A*02:01 were subject to linear fitting with R^2^ values shown. HLA-A*24:02 was excluded due to the limited range of data available. Non-binding peptide-HLA combinations that have no exact measured binding affinity value available were plotted at y=7. These data points were not included when performing linear regression. **b**, Measured binding categories plotted against predicted binding affinity values. **c**, Predicted binding affinity values plotted against average MFI values (measured at 100 nM concentration). **d**, Measured binding categories plotted against MFI average value (measured at 100 nM concentration).

We further examined how each prediction algorithm performed when compared to the measured IC50 binding values (**Data file S7; Fig. S10**). Out of the eight algorithms used, HLA-A*02:01 had the highest correlation scores for 6 out of the 8 algorithms, consistent with our observation that it has the largest training dataset (**Data file S3**) and thus was expected to yield the best accuracy. Overall, NetMHCpan came out on top, showing the highest correlation values in all 4 HLA alleles where sufficient data was available (**Fig. S11**). These experimental validation results combined help show the distinct anchor trends that exist among HLA alleles.

### Neoantigen prioritization results are influenced by accounting for anchor locations

Current pipelines fail to take into account HLA allele-dependent effects on anchor locations and researchers lack specific tools and databases to make use of such information. While the decision of whether a neoantigen should be prioritized over others involves many aspects not discussed in detail here (including variant allele frequencies, gene expression, and manufacturability considerations to name a few), we used a straightforward approach to evaluate the effects of introducing improved anchor information on neoantigen prioritization. Interpretation of a strong binding MT peptide candidate may depend on other factors such as WT binding affinity, agretopicity, mutation position, and anchor location(s), leading to different choices when prioritizing neoantigens (**Fig. 7a**). If the mutation is not at an anchor location, regardless of the WT peptide binding affinity, the MT peptide should be prioritized. In this case, the sequence for TCR recognition contains a mutation and will not be subject to central tolerance (**Fig. 7a**; scenario 1,2). Additionally, if the WT peptide is a weak binder, the MT peptide should be accepted regardless of whether the mutation is at an anchor location since both the MT and WT sequences have not previously been exposed to the immune system and therefore not subject to tolerance (**Fig. 7a**; scenario 4). However, if the WT binds strongly, regardless of agretopicity, and the mutation is at an anchor location, then this neoantigen will likely be subject to central tolerance and should be rejected from prioritization (**Fig. 7a**; scenario 3). These scenarios are considered when performing the following anchor position impact analysis.

**Fig. 7.**
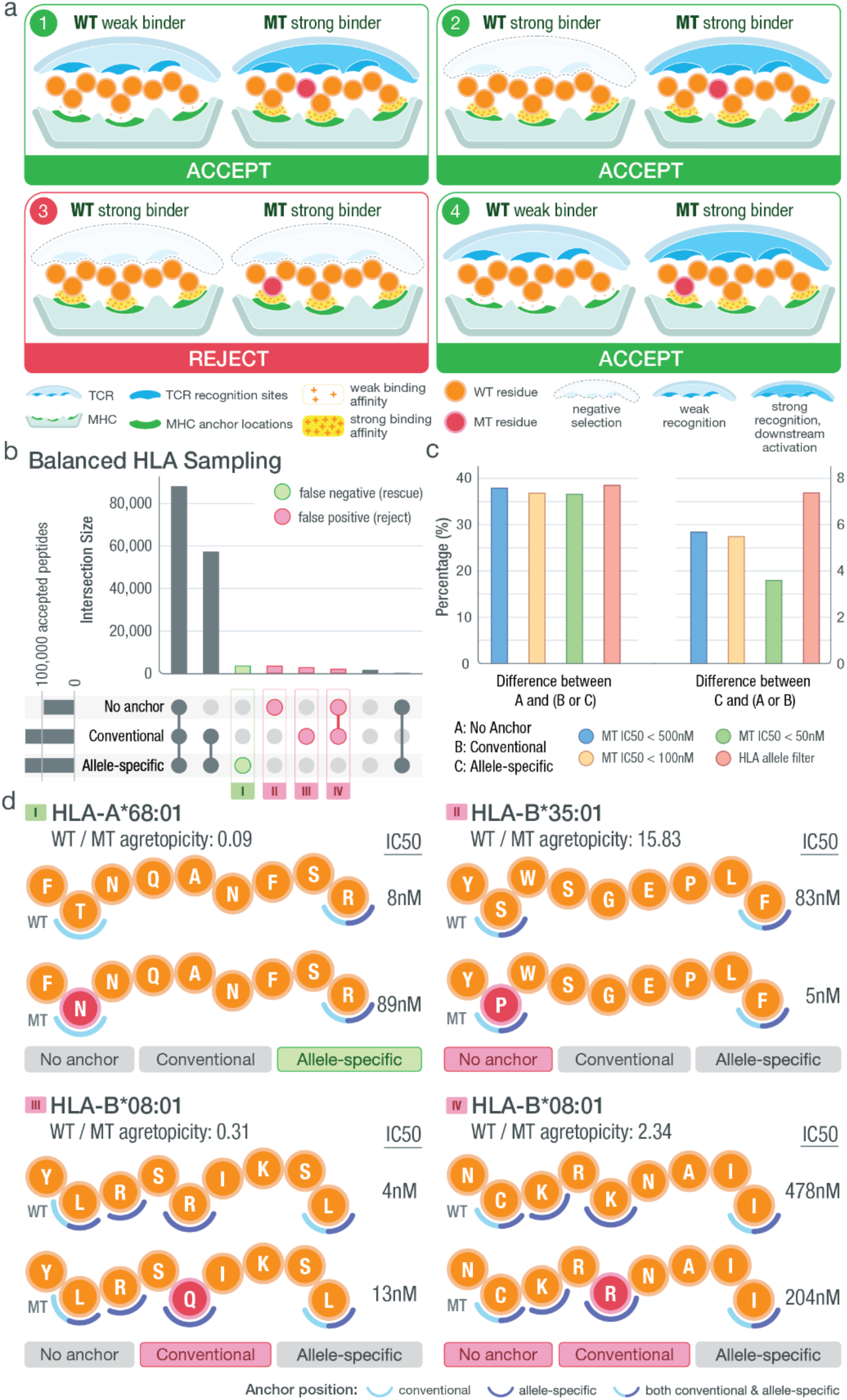
Impact of anchor position information on neoantigen prioritization decisions. **a,** Illustration of different scenarios that could be encountered when prioritizing neoantigens. Each circle represents a peptide residue with mutated residues marked in red. Anchor locations of the MHC are marked in green while TCR recognition sites are in blue. Predicted binding affinities of the MT/WT peptides are indicated using a yellow density field where higher density represents strong binding, and lower density represents weak binding. Three different scenarios of T-cell recognition are depicted. **b,** Upset plot showing the number of intersecting peptides based on those prioritized with no anchor filter (binding affinity < 500 nM and aggretopicity > 1), conventional filter (filtering based on conventional anchor assumptions) or allele-specific filter (filtering based on our computationally predicted anchor locations). Samples included for analysis were chosen such that HLA alleles were balanced appropriately. Peptides characterized differently between no anchor/conventional filter and allele-specific filter were categorized into false negatives (green circle) and false positives (red circle) with the assumption that the allele-specific filter produced more accurate results. **c,** Bar plot showing the percentage difference among accepted neoantigen candidates based on filters discussed in panel b. The filters in the legend represent how each starting dataset was filtered: 1) filtered by a strong binding cutoff of either 500 nM, 100 nM or 50 nM and 2) filtered based on the anchor patterns of the corresponding HLA allele. Our HLA allele filter excluded all peptides binding to HLA alleles with a canonical anchor pattern (e.g. [2,9] for 9-mer peptides). **d,** Examples of false positive and false negative peptides from each of the four subsets as marked in panel b. Matching HLA allele, peptide sequence, mutation position (red), median WT/MT IC50 values and fold changes are shown accordingly. Two sets of anchor locations are depicted for each scenario using semi-circles: conventional anchors are marked with light blue and allele-specific anchors are marked with dark blue. Positions where the two sets of anchors overlap are marked with split coloring of the semi-circle.

Our cohort impact analysis involved an additional set of TCGA patient samples where neoantigens were predicted for 923 selected patient-HLA allele pairings. When selecting patient-HLA paired samples we chose from a balanced HLA allele distribution (**Methods**). Our intent was to give a more balanced view of the impact across all HLA alleles without overt bias for the most common alleles. We first examined the proportion of SNV-induced neoantigens that went into each of the 4 scenarios described in **Fig. 7a** and found that 17%, 57%, 8% and 18% of all peptide candidates fell into scenarios 1, 2, 3 and 4 respectively (**Fig. S12**). All potential neoantigens were filtered according to three different criteria: A) mutant IC50 < 500 nM and agretopicity > 1 (‘no anchor filter’), B) supplementing this with a conventional anchor assumption (‘conventional filter’), or C) using our computationally predicted anchor locations (‘allele-specific filter’) (**Methods**). Peptide results showed that under the no anchor filter 57.9% of neoantigens are accepted compared to 93.2% under the conventional filter and 94.8% under the allele-specific filter, showing an overall net gain in the number of peptides when taking anchor considerations into account. When comparing filtered data sets under different criteria, 38.3% of neoantigens are potentially misclassified using the no anchor filter, and approximately 5.7% of candidates are potentially misclassified between the conventional filter and the allele-specific filter (**Fig. 7b**). These misclassifications involve the inclusion of peptides that are likely to be subject to tolerance (and could lead to false positives) and exclusion of peptides that could be strong candidates (false negatives). We repeated this analysis to see if our observations are consistent when 1) using different binding affinity cutoffs for strong-binding peptides (< 100 nM and < 50 nM) and 2) looking exclusively at peptide-MHC pairings that have a non-conventional anchor pattern (**Fig. 7c; Methods**). Our results showed that when comparing results between the no anchor (A) and any anchor considerations (B or C), differences of accepted candidates are similar across the four criteria (< 500 nM, < 100 nM, < 50 nM, and HLA filter). When comparing results between the allele-specific considerations (C) and the more naive approaches (A or B), we see an increase in percentage of difference with our HLA allele filtered dataset indicating a greater impact on neoantigen prioritization when using our allele-specific anchor method. This was expected since peptides analyzed in this dataset corresponded to HLA alleles with non-conventional anchor patterns. We also see a decrease in percentage when applying a binding affinity cutoff of 50 nM which we believe is in part due to less HLA alleles passing the stricter binding cutoffs (81 versus 98 unique HLA alleles found in datasets with cutoffs 50 nM and 500 nM respectively). By comparing peptides prioritized using the allele-specific anchor filter and those from the no anchor/conventional anchor filters, we highlighted the potential sources for false positives and false negatives (**Fig. 7b**). Examples of each scenario were pulled from our dataset to show how peptides passed or failed individual filters (**Fig. 7d**).

We additionally performed a patient-level impact analysis using 100 randomly selected TCGA samples, and predicted neoantigens each with their full set of class I HLA alleles (up to 6) (**Methods; Data file S8; Fig. S13a,b**). The neoantigen candidates were prioritized using the same set of criteria applied in the previous cohort analysis. We observed a significant impact on neoantigen prioritization results depending on the chosen filtering criteria. Specifically between the no anchor filter and the allele-specific filter, 99% of the patients analyzed had at least one neoantigen decision changed, with a median of 11 peptides with altered decisions, per patient. Similarly, between the no anchor filter and the conventional filter, 98% of the patients analyzed had at least one decision changed (median: 11 peptides) and between the conventional filter and the allele-specific filter, 65% of the patients had at least one decision changed (median: 1 peptide) (**Fig. S13c,d,e**). These results show the potential widespread effect of anchor considerations on patient-level prioritization results.

## Discussion

We developed a computational workflow for predicting probabilities of anchor positions for a wide range of the most common HLA alleles. Our results show that anchor positions vary substantially between different HLAs. A subset of our prediction results were confirmed by analyzing available crystallography structures of peptide-MHC complexes. We further experimentally validated HLA allele anchor patterns using binding assays and cell-based stabilization assays. The underlying quantitative scores from our anchor prediction workflow are available for incorporation into neoantigen prediction workflows and we believe this will improve their performance in predicting immunogenic tumor specific peptides. For simplicity, our illustrations have depicted peptide residues as either anchoring or potentially participating in TCR recognition, although previous research has shown that heteroclitic peptides can alter both simultaneously*(22)*. Hence, anchor residues and TCR recognition sites should not be considered mutually exclusive and should ideally be interpreted quantitatively where the anchor scores (provided in **Data file S2**.) reflect the probability of a peptide position participating in binding.

Using an independent pool of TCGA samples, previously excluded from the computational prediction process, we show that consideration of anchor prediction results can have a significant impact on neoantigen prioritization. In this study, the choices of whether to accept or reject a prioritization decision were based on hard cutoffs determined using an objective strategy across all HLA alleles. However, when making clinical decisions, we recommend using the actual anchor probabilities as guidance when prioritizing candidates. In most neoantigen characterization workflows, numerous other factors are taken into account to arrive at an overall prioritization decision, which may further increase differences between filtering strategies. Additionally, while our anchor results represent overall averaged scores across multiple neoantigens, slight variations of patterns do exist among different peptides for the same HLA allele. It would generally be computationally prohibitive to perform our workflow on a large scale for all possible neoantigen candidates. However, as a compromise, one could repeat our detailed process to generate peptide specific anchor predictions after performing other filtering strategies (e.g. genomic information including VAF and expression) and arriving at a shortlist of candidates.

Anchor results not only impact the selection process of neoantigens for personalized cancer vaccines, but also change the way neoantigen load estimation is currently defined. Neoantigen load estimation is commonly defined as the number of peptides whose binding affinity passes a certain threshold. However, this threshold, meant to limit to the approximate number of strong binding neoantigens, should also take into account the mutation position, HLA specific anchor locations, and agretopicity for more precise estimation. Our anchor impact analysis demonstrates the effect of this alteration on estimation results. Moreover, our analysis results show that there is a net gain of neoantigen candidates when taking anchor considerations into account compared to the commonly used agretopicity filters. This becomes important in the context of neoantigen prioritization, particularly when the minimum number of peptide vaccine candidates cannot be met for patients due to low tumor mutational burden. However, we also acknowledge that while our approach improves sensitivity it could come at the cost of slightly decreased specificity for patients with high neoantigen burden.

The neoantigen selection process requires careful consideration of numerous aspects, which have been discussed extensively*(6)*. In general, neoantigen-based vaccines act by stimulating the patient’s immune system for the production of activated cytotoxic T cells. However, compared to viral antigens where the protein sequence is entirely foreign, neoantigens, particularly those developed from SNVs, have merely subtle differences from the individual’s wildtype proteome. Thus, the need for a WT versus MT peptide comparison, while considering anchor locations, is an aspect specific to tumor neoantigens that other vaccine development pipelines could generally ignore. Though neoantigens derived from in-frame or frameshift indels diverge more from the WT sequence and are generally less influenced by our findings, cases where such mutations are located towards the beginning or end of a neoantigen may still cause anchor disruption in an allele-specific manner. Additionally, more work should be done to characterize the similarity of neoantigen candidates and the patient’s wildtype proteome for an overall accurate prioritization process.

Recent work, such as that from the Tumor Neoantigen Selection Alliance (TESLA)*(24)*, have hinted at the potential importance of anchor locations. In that study, researchers made the unexpected observation that among the 37 positively validated neoantigen candidates, none of the peptides had a mutation at position 2, a common anchor position for a range of HLA alleles, despite a high number of prioritized neoantigens with a position 2 mutation. Possible explanations could include the fact that neoantigens with the mutant residue at a strong anchor position have a disadvantage over those present at TCR sites as they require their WT counterpart to be a poor binder and the threshold for determining sufficiently weak binding of the WT peptide is unclear. However further investigation is required to address questions raised by such observations.

In addition to the limitations of being applicable to a subset of neoantigens derived from SNVs and certain indels, our work could be expanded to a wider range of HLA alleles. A larger HLA-peptide seed dataset could be achieved through a wide-scale prediction of strong binders for rare HLA alleles by mutating the wildtype proteome. There have also been publications noting the sequence motifs in HLA Class II binding peptides *(41)*, as well as others using molecular docking approaches *(42)* in an attempt to estimate anchor positions for HLA Class II alleles. However, class II alleles differ substantially from class I due to the difference in length of binding peptides and the binding pocket being more open and composed of a protein dimer. Additional data generation and further research will be required for the expansion of our workflow to address class II, though we hypothesize that the same principles should be applicable. Furthermore, while x-ray crystallography structures show support for our anchor location predictions, experimental validation with neoantigens designed to induce T-cell activation is needed to explicitly showcase the importance of our results in clinical settings. Although numerous clinical trials using neoantigen-based vaccines are underway, results published show a low accuracy for current neoantigen prediction pipelines*(43)*. By accounting for additional positional information, we hope to significantly reduce the number of false positive candidates and rescue false negative neoantigens to increase prediction accuracy. A prioritization strategy utilizing anchor results has been incorporated into the visual reporting of our neoantigen identification pipeline pVACseq*(7)*.

Machine learning algorithms have been widely applied in the context of neoantigen binding predictions. However, machine learning models trained on experimentally validated data with T-cell activation results are lacking and identifying features for these models is an active area of research. Anchor location probabilities may serve as an additional feature in machine learning model training on clinical data. This will allow for a more nuanced approach where anchor probabilities may be weighted for each allele-peptide pairing accordingly. These results and tools will help streamline the prioritization of candidates for neoantigen vaccines and may aid in the design of more effective cancer vaccines.

## Materials and Methods

### Study Design

The objective of this study was to predict allele-specific MHC anchor locations and investigate whether use of this information can significantly influence neoantigen identification and prioritization. We established a computational approach based on an ensemble of MHC binding prediction algorithms and exhaustive in silico mutation of peptides. By application of this approach to 609,807 distinct peptide sequences we were able to predict anchor location probabilities for 328 common HLA alleles with peptides of varying lengths (8-11 mers). Anchor location probabilities were clustered using hierarchical clustering and representative trends were identified and experimentally validated using various methods, including: x-ray crystallography structures, IC50 competition binding assays, and cell-based stabilization assays. All stabilization assays were completed 2-3 times in duplicate (N=4-6). Use of allele-specific anchor predictions was compared against conventional prioritization methods to demonstrate the potential influence on neoantigen prioritization results.

### Input data for identifying strong binding seed peptides for anchor site prediction

We assembled peptide data from various sources where binding prediction data were available through clinical collaborations and supplemented these with TCGA datasets where necessary to achieve better representation of less common HLA alleles. Datasets from clinical collaborations that were incorporated include 7 triple-negative breast cancer samples, 54 lymphoma samples, 20 glioblastoma samples, and 6 melanoma samples. Additionally, we mined data from 9,216 TCGA samples to optimize the number of strong binding peptides matched to each HLA allele by adding 10 samples for insufficient (<10 strong binding peptides) and 15 samples for previously unseen HLA alleles. Of these, 1,356 TCGA samples were used to generate seed HLA-peptide combinations to be used for downstream simulations. High quality variants included from TCGA samples were obtained from the Genomic Data Commons and selected according to their filter status (pass only) and required to be called by at least 2 out of 4 variant callers as previously described*(44)*. Peptide lengths considered ranged from 8- to 11-mers. In total, these datasets corresponded to 1,443 tumor samples, representing 328 matching HLA alleles, with 737 of these having more than 10 strong binding peptides, and a grand total of 609,807 strong binding peptides for use in the following analyses (**Fig. S1; Data file S1**).

### Computational prediction of anchor site positions for 328 HLA alleles

Peptides collected from input datasets were first filtered for strong binders using a binding affinity cutoff of 500 nM. These were used to build a seed dataset consisting of peptides predicted to be strong binders to individual HLA alleles. We first performed a saturation analysis to determine the appropriate number of random peptides needed to obtain a robust estimate of the likely anchor site locations of each HLA allele. This was done using peptides collected for HLA-A*02:01, where over 1,500 peptides were obtained for each peptide length. Random sampling with a subset size of 10 peptides showed consistently high correlation (> 0.95) with the largest subset size used where all 1000 peptides were incorporated (**Fig. S4**). Thus in downstream analysis, for each unique HLA and peptide length combination, 10 peptides were randomly selected from the database. For each of the 10 starting peptides at each position n, we obtained a score that reflects how much a mutation at this position (by mutation to all 19 other amino acid identities) will affect the overall binding affinity:

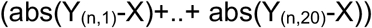

where X is the binding affinity of the unmutated peptide, and Y_(i,j)_ is the binding affinity of the peptide mutated at position i to amino acid number j (total of 20 possible amino acids to mutate to). All binding affinities were calculated using pVACbind from pVACtools (version 1.5.0)*(7)* in which the following algorithms were selected: NetMHC*(31)*, NetMHCpan*(32)*, MHCnuggets*(33)*, MHCflurry*(34)*, SMM*(35)*, Pickpocket*(36)*, SMMPMBEC*(37)*, NetMHCcons*(38)*. The median binding affinity across all 8 of these algorithms was used both to nominate strong binder peptides for the seed dataset, and to assess the impact of in silico mutation at each position of these peptides. For each strong binding peptide in the seed dataset, this systematic in silico mutation led to [length x amino acid identity x algorithm] binding predictions (e.g. 1,368 binding predictions for a single 9-mer peptide).

Each position was assigned a score based on how much binding affinity values were influenced by mutations at that position. These scores were then used to calculate the relative contribution of each position to the overall binding affinity of the peptide. Positions that together account for 80% of the relative overall binding affinity change were assigned as anchor locations for further impact analysis.

### Evaluating different seed peptide sources and their impact on the anchor patterns predicted

Three additional sources were explored and compared to peptides from our seed dataset to investigate whether the source of these strong binding peptides would influence our results. For comparison, we selected 1,000 strong binding peptides (median predicted binding affinity < 500nM) from the following four sources: randomly generated peptides, peptides randomly selected from the WT proteome, peptides randomly selected from a viral genome, and peptides from our seed dataset. To obtain randomly generated peptides, we first generated a random list of amino acids (100,000,000 aa in length) using a random choice function and tested 50,000 9-mer peptides to reach 1000 predicted strong binders. For peptides generated from the WT human proteome (ensembl) and the viral proteome (variola virus), we randomly selected 30,000 9-mer peptides from proteins with valid sequences longer than 9 amino acids.

For each strong binding peptide, we followed our computational workflow for anchor predictions and mutated each position of the peptide to all possible amino acids and accessed the change to binding respectively. Normalized scores across all peptide positions for 1,000 peptides were plotted in Fig S4.

### Bias analysis for HLA allele-specific anchor patterns

To evaluate whether our anchor patterns are influenced by the training data available for each HLA allele, we downloaded the training dataset of NetMHCpan4.0. Additionally, for HLA alleles with no training data available, we determined which HLA allele was its closest neighbor defined by NetMHCpan4.0 and what the distance measurements were between the neighboring alleles. With these data obtained, we used CytoScape to visualize the relationships between all alleles using network graphs. Each center node represents an HLA allele with training data (size of dataset correlates with the size of each node) and each connected node represents a neighboring HLA allele that uses the center node allele for its binding estimations. Distances are marked on the edges of each graph. Colors of each node reflect the anchor cluster assigned as shown in Fig 3.

### Input data for orthogonal evaluation of predicted anchor sites

To evaluate our anchor predictions, we collected 166 protein structures (pdb format) of peptide-MHC complexes and 61 peptide-MHC-TCR complexes from the Protein Data Bank*(45)* by querying for structures containing macromolecules matching class I HLAs. Structures were additionally reviewed to ensure valid peptide length (8–11) and those with TCRs attached were separated into a different list for downstream analysis to allow accurate solvent-accessible surface area (SASA) calculations. The HLA-peptide structures corresponded to 33 HLA alleles with peptides of varying lengths (8 to 11mer), while the HLA-peptide-TCR structures corresponded to 12 HLA alleles. A complete list of PDB ids selected for this analysis can be found in **Data file S4**.

### Orthogonal validation of predicted anchor sites by analysis of pMHC structures

The structures of peptide-MHC molecules were analyzed to infer potential anchor locations/residues. All PDB structures were analyzed in python using the MDTraj package*(46)*. For each position of a peptide bound to an HLA, we utilized two different metrics: 1) minimum distance of non-backbone atoms to all HLA associated atoms and 2) estimated solvent-accessible surface area (SASA) of the residue. In method 1, we calculated the distances between each atom of each residue and all HLA associated atoms. Non-backbone atoms were ordered by their distance to the closest HLA-associated atom and the top 50% were used to calculate an average distance representing an entire residue (with the exception of glycine where all atoms were considered). In method 2, we directly calculated the SASA of each residue (shrake_rupley function in MDTraj), which was used to infer the likelihood of being able to be recognized by the T-cell receptor. After calculating these values for each position of the peptide, they were directly compared to the anchor prediction scores by calculating a Spearman correlation. In the case of the distance metric, we expect positions of the peptide closer to the MHC to be more likely an anchor and those further (“bulging out”) to more likely interact with the TCR. Similarly, for the SASA metric, if a peptide position is more solvently accessible (higher SASA value) we expect it to be more accessible to the TCR as well and those that are less accessible would be more likely interacting with the MHC as an anchor.

For an overall evaluation of how well our anchor predictions correlated with these metrics (distance and SASA), Spearman correlations were determined across all structural data collected. For example, for a 9-mer peptide, a Spearman correlation was calculated for the 9 anchor prediction scores from the in silico mutation exercise compared to the 9 distance or SASA estimates obtained from the structure analysis. Out of 166 peptide-MHC structures collected, correlation values for 87 were plotted by randomly selecting at most 5 structures per HLA-length combination (**Data file S5**). For comparison, we also randomly shuffled distance and SASA scores across all positions of individual peptides and calculated correlation scores against this randomized dataset. The different sets of correlation values were then fit to Gaussian distributions (**Fig. 4d**). Non-paired two sample t-tests assuming unequal variance were performed to evaluate the differences among distributions (using python SciPy scipy.stats.ttest_ind).

Additional analysis was performed on the 61 peptide-MHC-TCR structures collected. After randomly selecting at most 5 structures per HLA-length combination, Spearman correlations derived from 31 structures were plotted. Correlations were calculated for 1) distance from peptide to HLA versus anchor prediction scores and 2) distance from peptide to TCR versus anchor prediction scores. Once again, the HLA-peptide distances were randomly shuffled and used as comparison and two sample t-tests were performed to evaluate the differences among distributions.

### Input data for evaluating the impact of anchor site considerations

To evaluate how anchor site considerations might influence neoantigen prioritization decisions, we considered a balanced HLA allele distribution when selecting input data. We randomly sampled up to 10 corresponding TCGA samples for each HLA allele with sufficient data (at least 3 out of 4 lengths have 10 or more matching peptides). 923 TCGA-HLA combinations were chosen from a total of 9,216 TCGA samples excluding the 1,356 used for the seed anchor site prediction data set described above. The 923 TCGA-HLA combinations corresponded to TCGA patients (**Data file S8**). To further evaluate impact of anchor considerations on a patientspecific level, an additional 100 TCGA patients were selected from the original 1,356 TCGA patient samples where we had neoantigen predictions for the patient’s full set of HLA alleles (**Data file S8**).

### Evaluating the impact of anchor site consideration on neoantigen prioritization

To analyze the importance of positional information on prioritization of neoantigens, TCGA patient samples were used as input and run through pVACtools (version 1.5.2) using the following options: -e 8,9,10,11, --iedb-retries 50, --downstream-sequence-length 500, --minimum-fold-change 0, --trna-cov 0, --tdna-vaf 0, --trna-vaf 0, --pass-only. The neoantigen candidates were then filtered and prioritized according to different criteria: **A**) Basic Filter: mutant peptide IC50 < 500 nM and agretopicity > 1, **B**) Decision based on a conventional anchor assumption that anchors are located at position 2 and the C-terminal position, **C**) Decision based on computationally predicted anchor locations. Specifically, under filter A) the position of a mutation with respect to MHC anchor positions is not considered. An accepted peptide means: MT peptide IC50 < 500 nM and WT IC50 / MT IC50 > 1, otherwise the peptide is rejected. Under filter B), anchor positions are defined to be 2 and n for all n-mer peptides. Under filter C), anchor positions are defined by our computational predictions, which are allelespecific. For both filters B and C, peptides are accepted if 1) the MT IC50 < 500 and WT IC50 > 500, or 2) the MT IC50 < 500, WT IC50 < 500 and the mutation is at a non-anchor location, as defined by the anchor definition of filters B and C respectively.

For filter C, anchor positions were defined individually for each peptide using the following strategy: Per anchor probability results from our computational workflow, each position of the n-mer peptide was assigned a score based on how binding to a certain HLA allele was influenced by mutations. These scores were then used to calculate the relative contribution of each position to the overall binding affinity of the peptide. Positions that together account for 80% of the overall binding affinity change were assigned as anchor locations for impact analysis. Assuming P_n_ represents the normalized probability of position n within the peptide, for each HLA allele, the anchor(s) is determined as following:

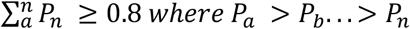

Filtered lists were then compared for overlap and differences. We also followed the same evaluation process for different starting candidate lists of varying stringency, including: candidates filtered with a strong-binding cutoff of 100 nM, candidates filtered with a strong-binding cutoff of 50 nM and candidates filtered by their binding HLA allele’s anchor patterns. For our HLA allele filtered dataset, all HLA alleles with an exact [2, n] anchor pattern (for n-mer peptides) were considered canonical and excluded from further evaluation.

For our cohort analysis, all neoantigen candidates were considered with no additional filtering. For our patient-level analysis, neoantigen candidates were processed additionally using the top_score_filter (“pVACseq top_score_filter” command of pVACtools) to generate top neoantigen candidates for individual variants. These top candidates were compiled and the same filters A, B, and C were used to determine prioritization decisions. The percentage differences between filters were calculated based on decisions for all top candidates for each individual patient.

### HLA coverage and population frequency

Global HLA allele frequencies were generated using data from the Allele Frequency Net Database*(39)*. The database contains HLA genotype data for Class I alleles across 197 distinct populations. Two populations in the database (“Chile Santiago” and “Russia Karelia”) did not have ambiguity-resolved HLA genotype data and were excluded from this analysis. Global HLA allele frequencies were calculated by (1) aggregating all 195 sample populations, (2) summing HLA allele counts over all sample populations, and (3) dividing HLA allele counts over total counts of the HLA gene across all populations. It should be noted that the HLA frequencies calculated do not reflect true global HLA frequencies since true population/region sizes were not considered. To calculate the percentage of population that our 328 HLA alleles affect, Class I alleles were split into respective subclasses of HLA-A, HLA-B and HLA-C. Global frequencies were summed in each subclass to obtain the percentage of population potentially affected by our HLA allele anchor results.

### Selection of validation peptides

In order to select a group of peptide-HLA combinations for validation experiments, we first examined our seed database of predicted strong binding combinations of peptides and HLA alleles. The database was then filtered for preselected HLA alleles with varying anchor patterns.

For each HLA allele within an anchor subgroup, we then selected 3-5 peptides and performed validation experiments to measure their binding affinity. Validated peptides were then mutated at various positions (including those predicted to be anchors and non-anchors) to evaluate amino acid changes and their effect on binding affinity. Due to the limited resources and high cost of synthesizing peptides and performing validation experiments, we strategically chose 4 different amino acid mutations for each position evaluated. Both positions and amino acid mutations were chosen to optimize informativeness. The amino acid mutations were selected based on their predicted influence on binding affinities by two methods. For our first batch of peptides and their in-depth analysis, we selected two of the most/least disturbing mutations each according to our predictions. For our second batch of peptides, this strategy was further optimized to select 1 mutation from each quartile based on predictions of how much they disturb the binding interaction. These mutated peptides were then synthesized by GenScript (Piscataway, NJ, USA) for further validation. The peptides were ordered with the following specifications: quantity: 4 mg, weight: gross, purity: ≥75%, delivery format: lyophilized, aliquoting to vials: 4.

### Peptide:MHC IC50 binding assays

As previously described*(47)*, peptide binding validation assays to determine IC50 values were carried out by Pure Protein LLC, utilizing the method of fluorescence polarization (FP). The technique is unique among methods used to analyze molecular binding events because it allows the instantaneous measurement of the ratio between free and bound labeled ligands in solution without any separation steps. FP is based on the principle that if a fluorescent-labeled peptide binds to the soluble HLA molecule of higher molecular weight, polarization values will increase due to the slower molecular rotation of the bound probe. In this competition assay, a reference fluorescent-labeled peptide is incubated with activated sHLA in the presence of the neoantigen-derived peptide competitor and peptide/HLA interaction is monitored over time. A positive response occurs when the peptide of interest outcompetes the labeled peptide tracer. A negative response will take place when the peptide of interest has no binding characteristics and only the tracer is assembling with the sHLA. Competition experiments were analyzed by plotting FP values as a function of the logarithms of competitor concentrations. The binding affinity of each competitor peptide was expressed as the concentration that inhibits 50% binding of the FITC-labeled reference peptide. Observed half-maximal inhibitory concentrations (IC_50_) were determined by nonlinear curve fitting to a dose-response model with a variable slope using the specific software Prism (Graph Pad Software, Inc., San Diego, CA). In order to prioritize epitopes with greatest potential, ranked peptide IC_50_ values were classified as log-dependent values (x) based on preset affinity categories as described in Buchli et al. 2005*(47)*, into high (x< 3.7), medium (3.7 < x < 4.7), low (4.7 < x < 5.5), very low (5.5 < x < 6.0) and no binder (x > 6.0). Binding assays were performed for all (or part of) the peptides with the following matching HLA alleles: HLA-B*07:02, HLA-B*08:01, HLA-A*02:01, HLA-A*24:02, and HLA-A*68:01.

### Cell-based peptide:MHC stabilization assays

The recipient of all HLAs of interest in this study was a TAP deficient, class I negative cell line, that was created by introduction of HSV-ICP-47 (gift from Ted Hansen) using the pMIP (puro^r^) retrovirus and selection of puromycin resistant cells into the Class I negative, lymphoblastoid cell line, 721.221 (gift from M. Colonna, Washington University Saint Louis) cultured in Iscoves MEM, 10% (v/v) FBS with 0.6 ug/mL puromycin. HLA cDNA (IDP-IMGT/HLA) was prepared synthetically (Blue Heron, Bothell, WA) and shuttled into pMIG (GFP) retroviral vector. All HLA expressing cells were enriched by sorting GFP positive cells to greater than 95% using a Sony Synergy Flow Cell Sorter. For peptide stabilization assays, the cell line expressing the HLA, was washed twice in PBS and then serum starved for 1 hour in RPMI1640, 10 mM HEPES, 1x nonessential amino acids, 1 mM sodium pyruvate, 2 mM glutamine, prior to plating, washed twice and incubated with 100 ug/mL peptide of interest previously solubilized in 10% DMSO and sterile filtered through a 0.2 μm centrex filter. The cell suspensions were maintained at ambient temperature for 1.5-3 hours and then shifted to 37°C overnight at 5% CO2. To quantitate peptide stabilized complexes, the cell suspensions were then washed twice to remove unbound peptide and then incubated with APC conjugated W6/32 Mab (Biolegend, 311410) for 30 minutes at ambient temperature. Cells were washed twice and complexes were detected in the presence of 7-AAD to remove dead cells from the analysis. The median fluorescence intensity of 7-AAD negative, GFP/APC positive cells were quantitated by FlowJo v.10.6.1 after collection on a Beckman Coulter Navios three laser flow cytometer. Each HLA tested utilized a Flu viral positive control peptide known to stabilize the HLA of interest (**Data file S6**) and a negative control peptide that stabilized a different class I molecule. Each assay was completed 2-3 times in duplicate (N=4-6) and graphed in GraphPad Prism v.9.2.0 after subtraction of the MedFI of the no peptide control. The corrected mean fluorescence intensity (MFI) values are included in Data file S6. Stabilization assays were performed for all of the peptides with the following matching HLA alleles: HLA-A*02:01, HLA-A*23:01, HLA-A*24:02, HLA-A*31:01, HLA-A*68:01, HLA-B*07:02, and HLA-B*18:01.

### Statistical Analysis

Statistical analysis was performed using the python SciPy package. Non-paired two sample t-tests assuming unequal variance were performed to evaluate the differences among distributions for Fig. 4d. T-tests were performed using the scipy.stats.ttest_ind function. The normal distribution of variables was calculated using the Shapiro-Wilk test.

## List of Supplementary Materials

**Fig. S1**. Distribution of peptides collected per allele across 328 HLA alleles, split by peptide length

**Fig. S2**. Saturation analysis for evaluating subsample size of peptides needed for simulation analysis

**Fig. S3**. Hierarchical clustering of anchor prediction scores across all 8, 10, and 11-mer peptides assembled

**Fig. S4.** Comparison of anchor pattern across different seed peptide sources using HLA-A*02:01

**Fig. S5.** Analysis of potential for supporting algorithm bias across 328 HLA alleles

**Fig. S6.** Analysis of crystallography data for HLA-B*08:01 and 9-mer peptides

**Fig. S7.** Distribution of Spearman correlation values comparing anchor predictions and peptide-HLA/TCR distance measurements

**Fig. S8.** Additional experimental validation data for the predicted HLA-B*08:01 anchor pattern

**Fig. S9.** Additional experimental validation data for HLA-A*23:01 and HLA-A*68:01

**Fig. S10.** Breakdown of predicted binding affinity values versus measured binding affinity by individual algorithms

**Fig. S11.** Correlation scores between individual MHC binding algorithm predictions and measured binding affinities across HLA alleles

**Fig. S12**. Distribution of anchor scenario categories for neoantigen candidates from 923 tumor-HLA paired samples

**Fig. S13**. Patient-level analysis for impact of anchor considerations on neoantigen prioritization

**Data file S1**. Seed dataset of strong binding neoantigen candidates

**Data file S2**. Anchor predictions for 328 HLA alleles

**Data file S3**. HLA Summary Information

**Data file S4**. HLA PDB data table

**Data file S5.** Orthogonal validation correlation data

**Data file S6.** Summary of all in vitro and cell based experimental validation data

**Data file S7**. Breakdown of individual algorithm predictions and their correlation with validation data

**Data file S8:** List of TCGA samples for impact analysis

**Movie S1**. Demonstration of orthogonal validation using distance and SASA metrics from x-ray crystallography structures

## Acknowledgements

We thank the patients and their families for the donation of their samples and participation in clinical trials. The results shown here are in part based upon data generated by the TCGA Research Network: https://www.cancer.gov/tcga.

## Author contributions

H.X. and M.G.contributed to the design of the model and the computational framework. H.X. and S.S. contributed to the analysis of the data. M.B., O.C.O., R.B., E.M., P.P. designed and performed experiments and analyzed data. M.M.R., A.B., C.A.R., C.P., K.C.C., G.P.D., T.A.F., A.R., and W.E.G. contributed to acquisition and generation of patient sample data. H.X., J.H., S.K., S.P.G., T.M.J., C.A.M., W.E.G., O.L.G., M.G. contributed to study design. M.G. and O.L.G. contributed to the overall supervision of the project. H.X. and J.M. wrote the manuscript and created all figures in consultation with M.G. and O.L.G. All authors provided critical feedback and helped shape the research, analysis and manuscript.

## Data and materials availability

All data associated with this study are available in the main text or the supplementary materials.

