## Supplemental Figures for "Computational prediction of MHC anchor locations guide neoantigen identification and prioritization"

### Supplementary Figures

**Fig. S1. Distribution of peptides collected per allele across 328 HLA alleles, split by peptide length**

Histograms summarizing the distribution of predicted strong binding peptides collected for 328 HLA alleles. Peptides are plotted according to their respective lengths. The total number of peptides for each k-mer is shown as N. The x-axis represents the total number (X) of k-mer peptides for individual HLA alleles while the y-axis represents the number of HLA alleles with X number of peptides matched.

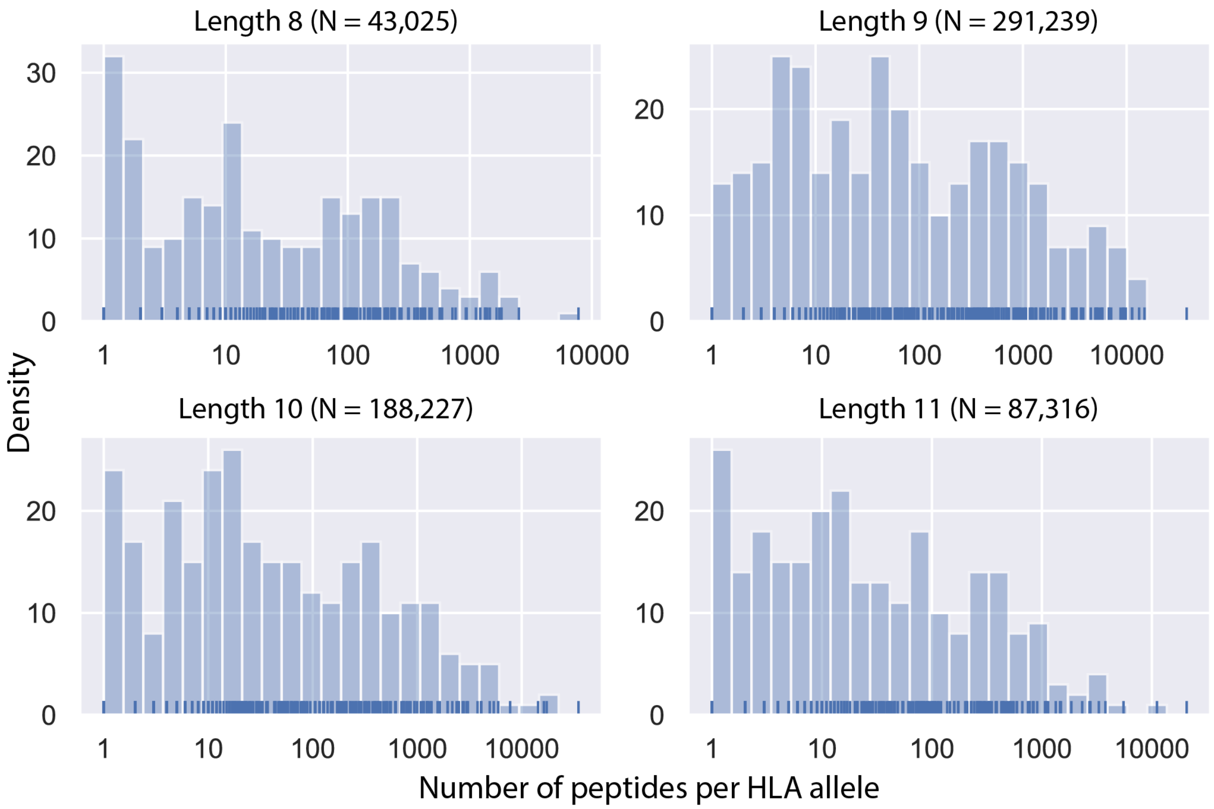

**Fig. S2. Saturation analysis for evaluating subsample size of peptides needed for simulation analysis**

Results of saturation analyses performed for 8-11-mer peptides using Spearman correlation are depicted as line plots with colors indicating random iterations. For each k-mer, 5 random rounds were performed with subset sizes of 1, 5, 10, 20, 50, 100, 200, 500 and 1,000. Correlations were calculated between each subset size and the largest subset (N=1,000) with the assumption that 1,000 samples was adequate to represent a ground truth for this purpose.

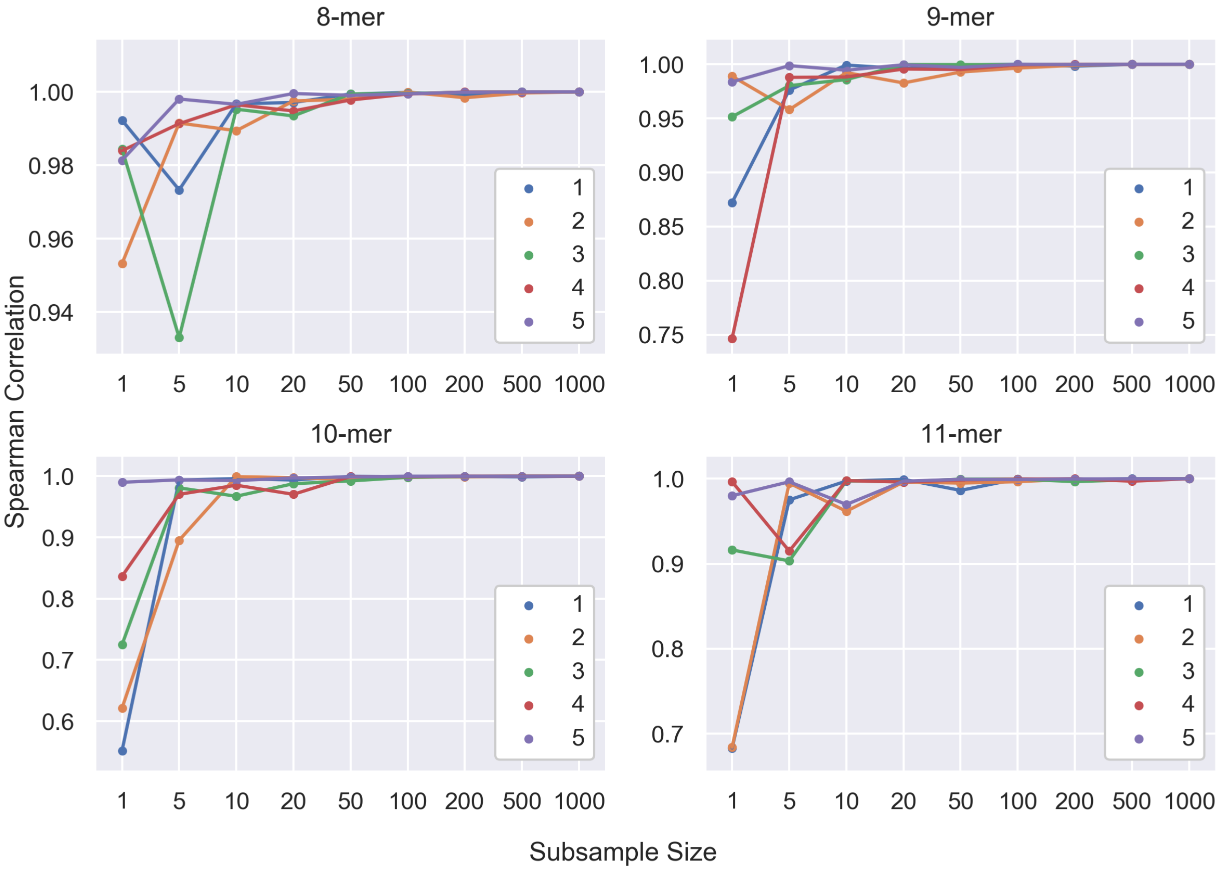

**Fig. S3. Hierarchical clustering of anchor prediction scores across all 8, 10, and 11-mer peptides assembled**

Heatmaps depict anchor prediction scores clustered using hierarchical clustering with average linkage across all 328 HLA alleles for which 8-mer, 10-mer and 11-mer peptide data were collected. For the individual heatmaps, the x-axis represents the k peptide positions and the y-axis represents the HLA alleles for which k-mer peptides were available.

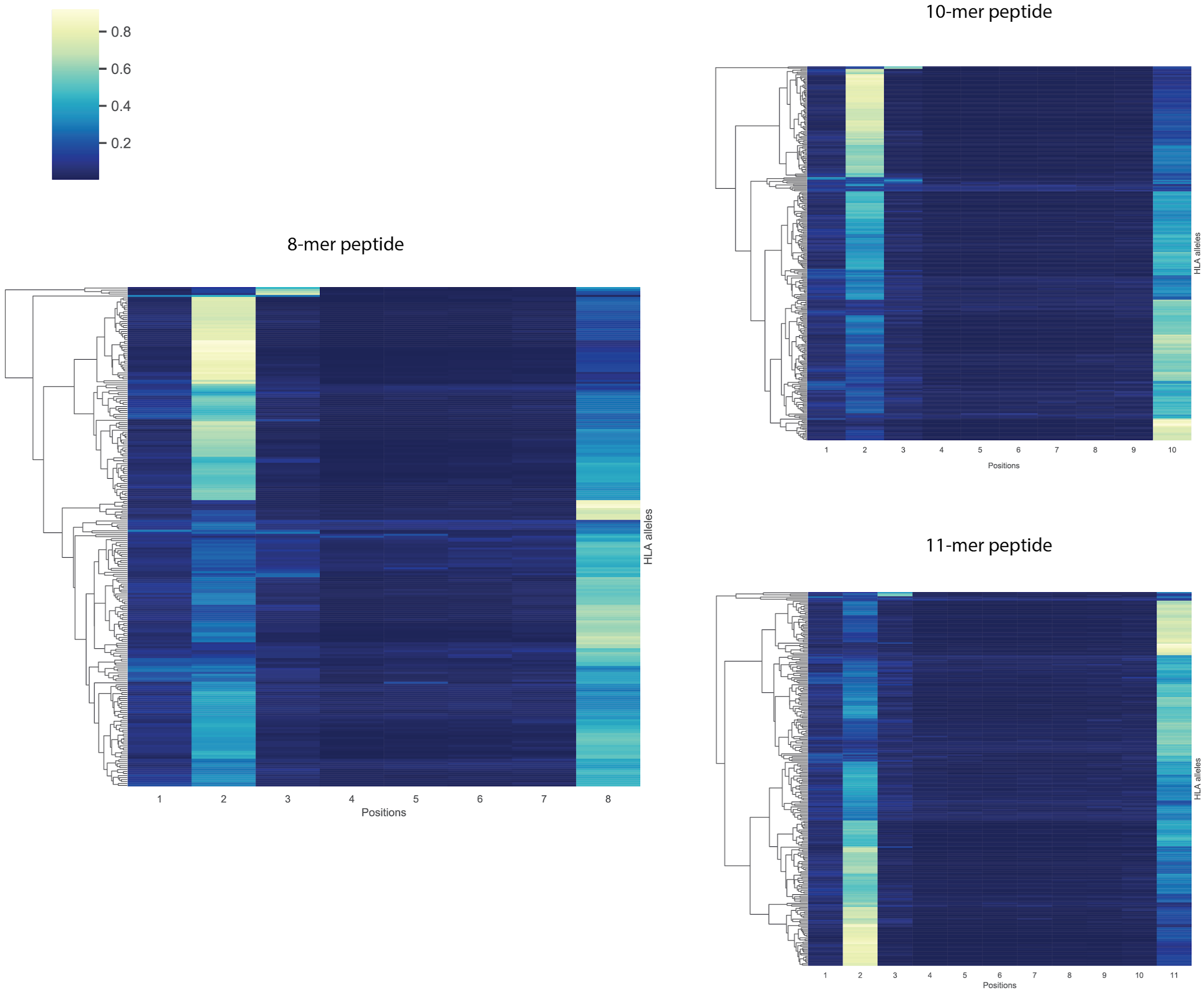

**Fig. S4. Comparison of anchor pattern across different seed peptide sources using HLA-A*02:01**

Anchor pattern for HLA-A*02:01 calculated from 1,000 peptides for each individual peptide source. We compared results from our seed dataset of peptides (tumor mutation derived) to three qualitatively distinct additional sources of peptides: 1) 9-mer peptides generated from a random 100,000,000 length sequence of amino acids, 2) random 9-mer peptides sampled from the human reference proteome (based on known Ensembl protein sequences) and 3) random 9-mer peptides from a viral proteome (variola). From each source, we identified 1,000 peptides that were predicted to be strong binders (Methods) and mutated each position for each amino acid following our computational workflow. We performed in silico predictions for each of the mutated peptides and calculated anchor probabilities based on all peptides collected. Normalized anchor probabilities are plotted on the y-axis and the peptide positions are listed on the x-axis.

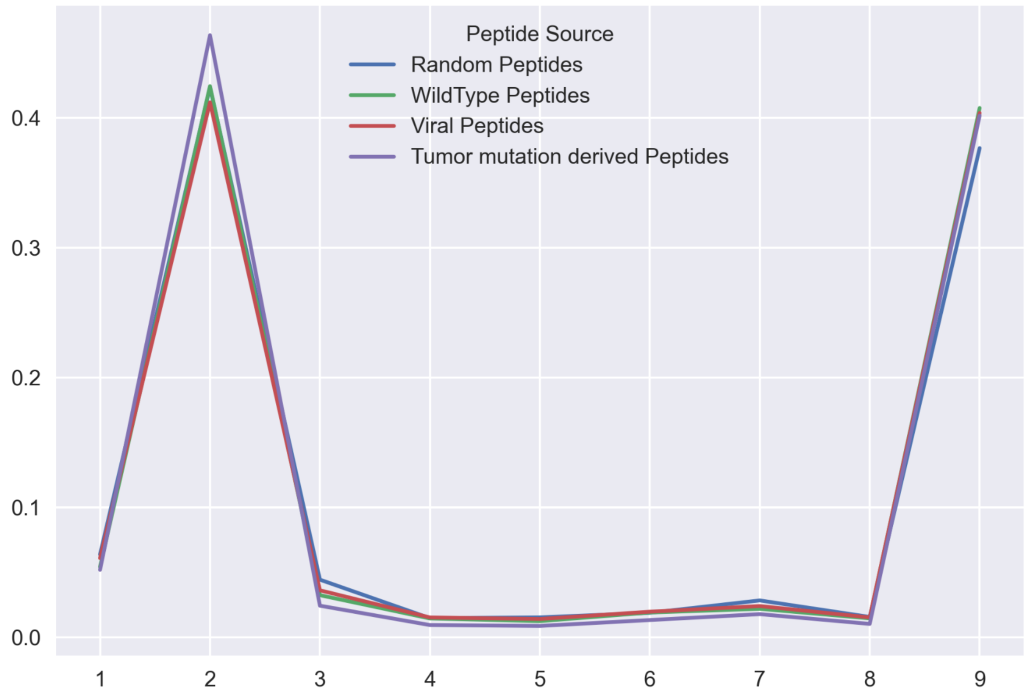

**Fig. S5. Analysis of potential for supporting algorithm bias across 328 HLA alleles**
**a,** Bar plot showcasing the number of HLA alleles being supported by the algorithms used. A maximum of 8 algorithms are available with the majority of the HLA alleles being supported by at least 4 algorithms. The specific breakdown is as following: out of the 328 HLA alleles, 54 HLA alleles are supported by all 8 algorithms, 2 are supported by 7 algorithms, 11 are supported by 6 algorithms, 8 are supported by 5 algorithms, 252 are supported by 4 algorithms and 1 is supported by 1 algorithm. **b,** Scatter plot showing distribution of HLA allele anchor pattern clusters with respect to two metrics: Mean value of variances calculated based on up to 8 different algorithmic predictions (x-axis) and distance to nearest HLA allele neighbor according to NetMHCpan (y-axis). If the HLA allele has training data available, then the distance is 0 since the nearest neighbor is only used in cases where the HLA allele in query needs to be estimated based on other similar HLA alleles. Colors are used to denote the clusters as annotated in Fig. 3. **c,** Violin plot showing the distribution of mean value of variances calculated across (up to 8) algorithms across the different anchor clusters. Lower variance indicates a better prediction consistency across the different algorithms used. **d,** Violin plot showing the distribution of distances across the different anchor clusters. Distance of 0 indicates that the HLA allele either did not need a closest neighbor to estimate binding or that the HLA pseudo-sequences between the two alleles were identical. **e,** Scatter plot showing the correlation of distance to the mean variance across different HLA alleles. **f,** Overview of a network graph showing how HLA alleles in our dataset are connected to each other. Each center node has training data available with the size proportional to the size of the center node. Edges connect nodes representing neighboring HLA alleles (as defined by NetMHCpan4.0). Color of each node represents the anchor cluster assigned as in Fig. 3. **g,** Zoomed in view of the network graph in **f** for the top 10 largest networks. Weights on edges reflect the distance between the HLA alleles (output from NetMHCpan4.0). Center HLA allele is listed on the top left of each box in bold.

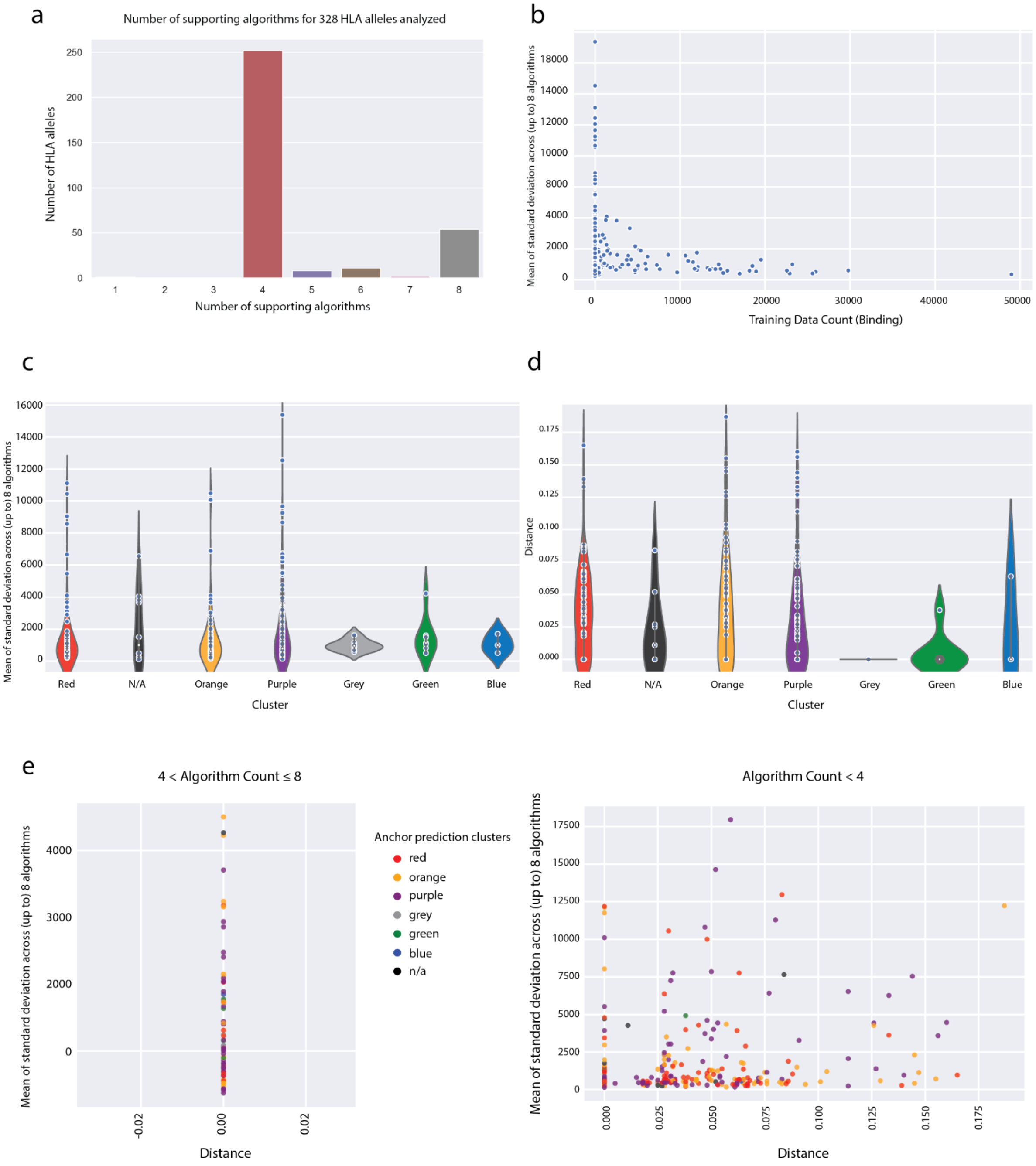
a
Number of supporting algorithms for 328 HLA alleles analyzed
Number of HLA alleles
Number of supporting algorithms
b
Mean of standard deviation across (up to) 8 algorithms
Training Data Count (Binding)
c
Mean of standard deviation across (up to) 8 algorithms
Red
N/A
Orange
Purple
Grey
Green
Blue
Cluster
d
Distance
Red
N/A
Orange
Purple
Grey
Green
Blue
Cluster
e
4 < Algorithm Count ≤ 8
Mean of standard deviation across (up to) 8 algorithms
Distance
Anchor prediction clusters
red
orange
purple
grey
green
blue
n/a
Algorithm Count < 4
Mean of standard deviation across (up to) 8 algorithms
Distance

f

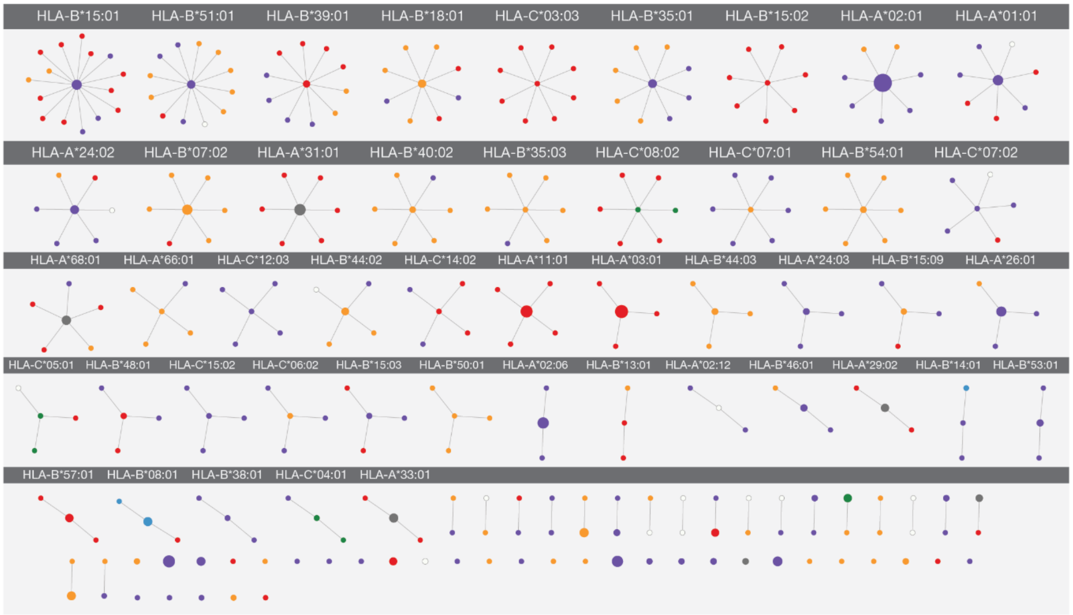

HLA-B*15:01
HLA-B*51:01
HLA-B*39:01
HLA-B*18:01
HLA-C*03:03
HLA-B*35:01
HLA-B*15:02
HLA-A*02:01
HLA-A*01:01
HLA-A*24:02
HLA-B*07:02
HLA-A*31:01
HLA-B*40:02
HLA-B*35:03
HLA-C*08:02
HLA-C*07:01
HLA-B*54:01
HLA-C*07:02
HLA-A*68:01
HLA-A*66:01
HLA-C*12:03
HLA-B*44:02
HLA-C*14:02
HLA-A*11:01
HLA-A*03:01
HLA-B*44:03
HLA-A*24:03
HLA-B*15:09
HLA-A*26:01
HLA-C*05:01
HLA-B*48:01
HLA-C*15:02
HLA-C*06:02
HLA-B*15:03
HLA-B*50:01
HLA-A*02:06
HLA-B*13:01
HLA-A*02:12
HLA-B*46:01
HLA-A*29:02
HLA-B*14:01
HLA-B*53:01
HLA-B*57:01
HLA-B*08:01
HLA-B*38:01
HLA-C*04:01
HLA-A*33:01


g

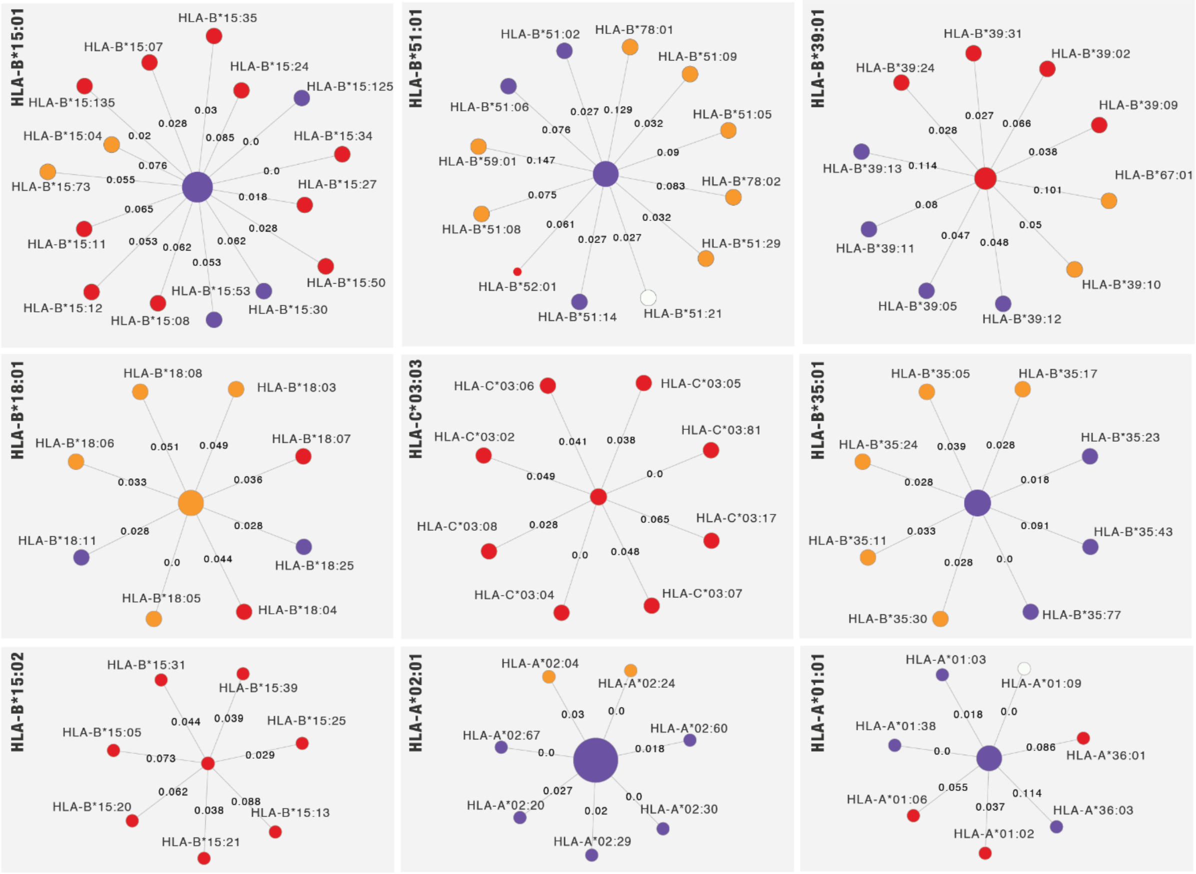

HLA-B*15:01
HLA-B*15:35
HLA-B*15:07
HLA-B*15:24
HLA-B*15:125
HLA-B*15:135
HLA-B*15:04
HLA-B*15:34
HLA-B*15:73
HLA-B*15:27
HLA-B*15:11
HLA-B*15:50
HLA-B*15:53
HLA-B*15:12
HLA-B*15:08
HLA-B*15:30
0.028
0.03
0.02
0.085
0.0
0.076
0.0
0.055
0.018
0.065
0.028
0.053
0.062
0.062
0.053
HLA-B*51:01
HLA-B*51:02
HLA-B*78:01
HLA-B*51:09
HLA-B*51:06
HLA-B*51:05
HLA-B*59:01
HLA-B*78:02
HLA-B*51:08
HLA-B*51:29
HLA-B*52:01
HLA-B*51:14
HLA-B*51:21
0.027
0.129
0.076
0.032
0.147
0.09
0.075
0.083
0.061
0.032
0.027
0.027
HLA-B*39:01
HLA-B*39:31
HLA-B*39:24
HLA-B*39:02
HLA-B*39:09
HLA-B*39:13
HLA-B*67:01
HLA-B*39:11
HLA-B*39:10
HLA-B*39:05
HLA-B*39:12
0.028
0.027
0.066
0.038
0.114
0.101
0.08
0.05
0.047
0.048
HLA-B*18:01
HLA-B*18:08
HLA-B*18:03
HLA-B*18:06
HLA-B*18:07
HLA-B*18:11
HLA-B*18:25
HLA-B*18:05
HLA-B*18:04
0.051
0.049
0.033
0.036
0.028
0.028
0.0
0.044
HLA-C*03:03
HLA-C*03:06
HLA-C*03:05
HLA-C*03:02
HLA-C*03:81
HLA-C*03:08
HLA-C*03:17
HLA-C*03:04
HLA-C*03:07
0.041
0.038
0.049
0.0
0.028
0.065
0.0
0.048
HLA-B*35:01
HLA-B*35:05
HLA-B*35:17
HLA-B*35:24
HLA-B*35:23
HLA-B*35:11
HLA-B*35:43
HLA-B*35:30
HLA-B*35:77
0.039
0.028
0.028
0.018
0.033
0.091
0.028
0.0
HLA-B*15:02
HLA-B*15:31
HLA-B*15:39
HLA-B*15:05
HLA-B*15:25
HLA-B*15:20
HLA-B*15:13
HLA-B*15:21
0.044
0.039
0.073
0.029
0.062
0.088
0.038
HLA-A*02:01
HLA-A*02:04
HLA-A*02:24
HLA-A*02:67
HLA-A*02:60
HLA-A*02:20
HLA-A*02:30
HLA-A*02:29
0.03
0.0
0.0
0.018
0.027
0.0
0.02
HLA-A*01:01
HLA-A*01:03
HLA-A*01:09
HLA-A*01:38
HLA-A*36:01
HLA-A*01:06
HLA-A*36:03
HLA-A*01:02
0.018
0.0
0.0
0.086
0.055
0.114
0.037

**Fig. S6. Analysis of crystallography data for HLA-B*08:01 and 9-mer peptides**
Results of two structures produced for HLA-B*08:01 with 9-mer peptides (blue and green lines). Top panel corresponds to distance measurements for each position while the bottom panel corresponds to SASA measurements. X-axis represents positions 1 to 9 of the peptides included.

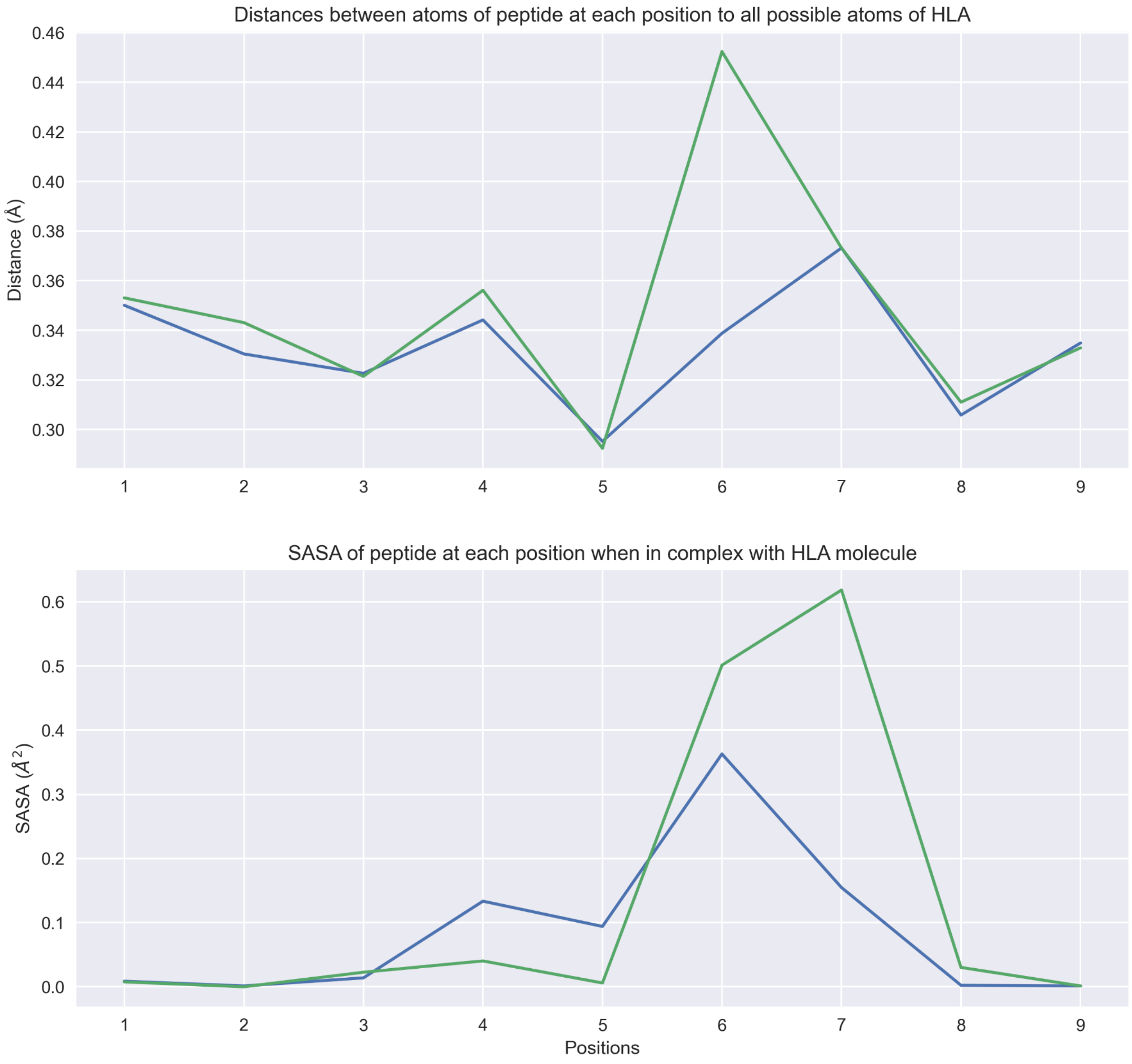

**Fig. S7. Distribution of Spearman correlation values comparing anchor predictions and peptide-HLA/TCR distance measurements**

This analysis is based on a collection of 61 structures obtained from PDB where a specific peptide, MHC and TCR were crystallized as a three part complex. The distribution of Spearman correlation values calculated by comparing our anchor scores to distance metrics obtained from these structures is shown. The blue line represents correlations between the predicted anchor score and the HLA-peptide distance measurements (expected to be negatively correlated because a position with a strong/high anchor score will tend to be closer to the MHC groove). The red line represents correlations between the predicted anchor score and the TCR-peptide distance measurements (expected to be positively correlated because positions acting as an MHC anchor will tend to be further away from the TCR interface). The green line represents the correlations between the predicted anchor score and randomized distances.

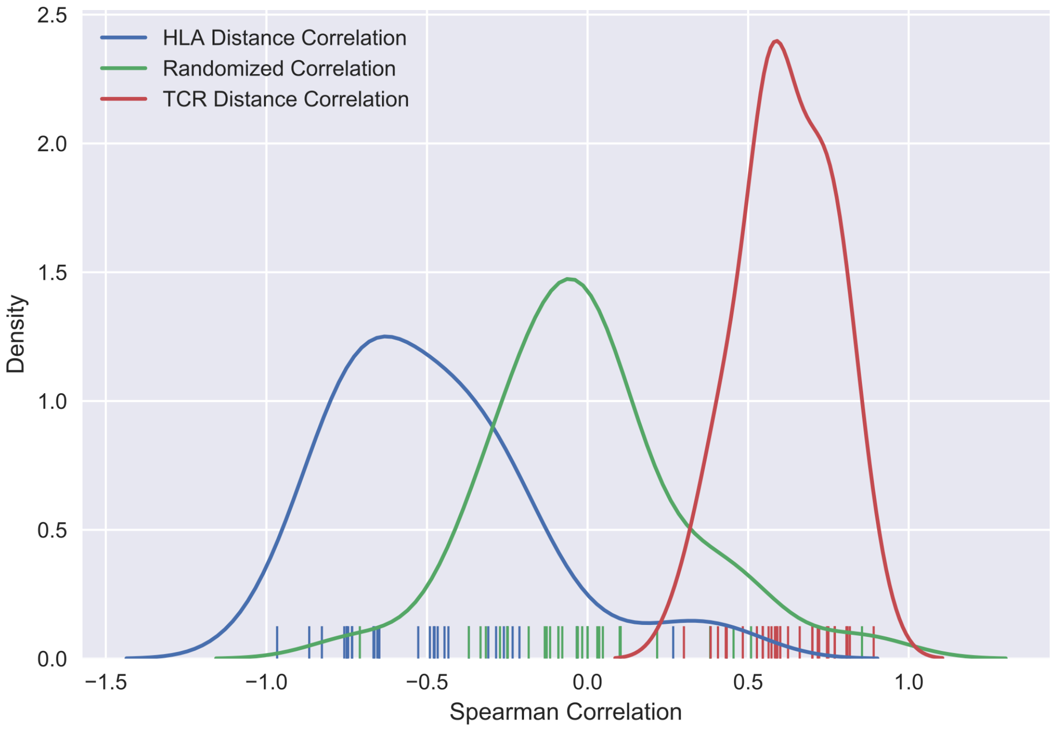

**Fig. S8. Additional experimental validation data for the predicted HLA-B*08:01 anchor pattern**

Validation results for HLA-B*08:01 with a predicted 2W-3W-5W-9M anchor pattern using peptide TLFMREHNL. **a**, Overall anchor prediction scores for each position of 9-mer peptides when binding to HLA-B*08:01. **b**, Binding affinity changes (y-axis) plotted for each position of the specific 9-mer peptide TLFMREHNL. Each position of peptide was mutated to 19 other amino acids to evaluate influence on binding affinity. **c**, IC50 values measured from binding affinity assays are plotted for both unmutated and mutated peptides. Peptides are marked by their mutation positions (P2, P3, P5, P6 and P9), predicted binding affinity values. Lower IC50 values correspond to stronger binding between the peptide and HLA allele.

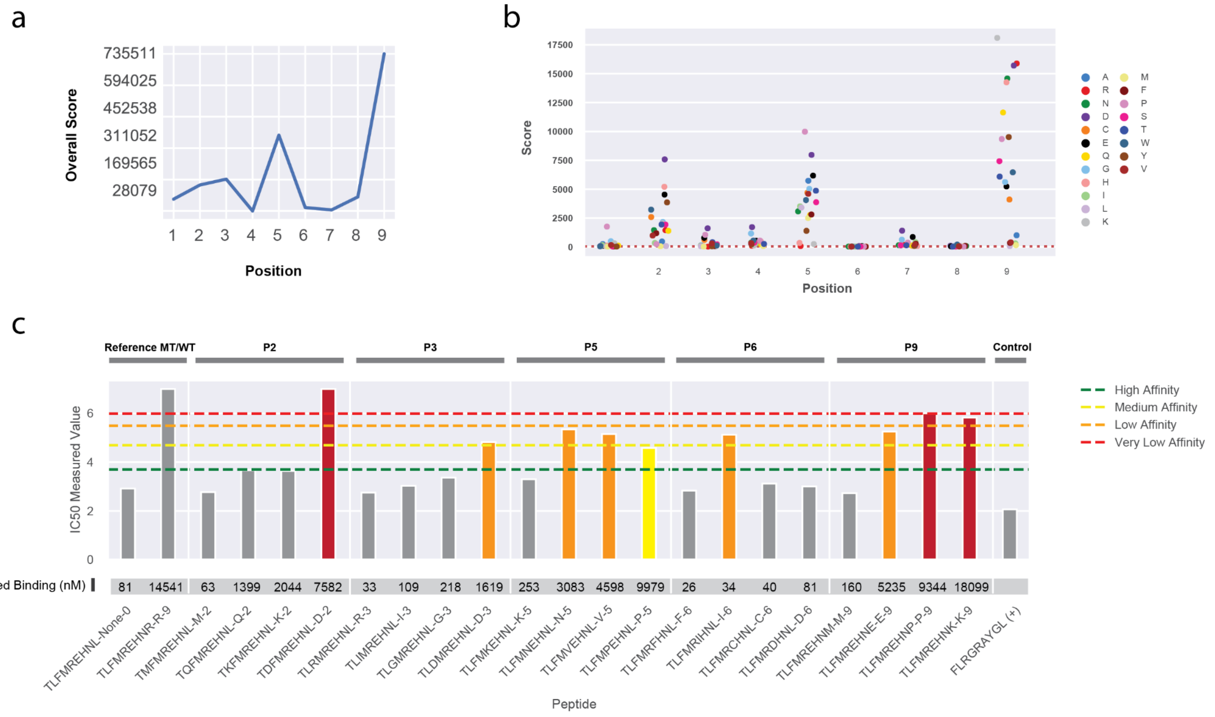

**Fig. S9. Additional experimental validation data for HLA-A*23:01 and HLA-A*68:01**
**a**, Validation results for HLA-A*68:01 with a predicted 9S anchor pattern using peptide ELAKHACPR. MFI values measured from cell stabilization assays are plotted for both unmutated and mutated peptides at 100 nM concentration. **b**, Validation results for HLA-A*23:01 with a predicted 2W-9S anchor pattern using peptide QWLQPEAHF. MFI values measured from cell stabilization assays are plotted for both unmutated and mutated peptides at 20 nM concentration. Peptides are marked by their mutation positions (P1, P2, P5, P9), and predicted binding affinity values. Higher MFI values correspond to stronger binding between the peptide and HLA allele.

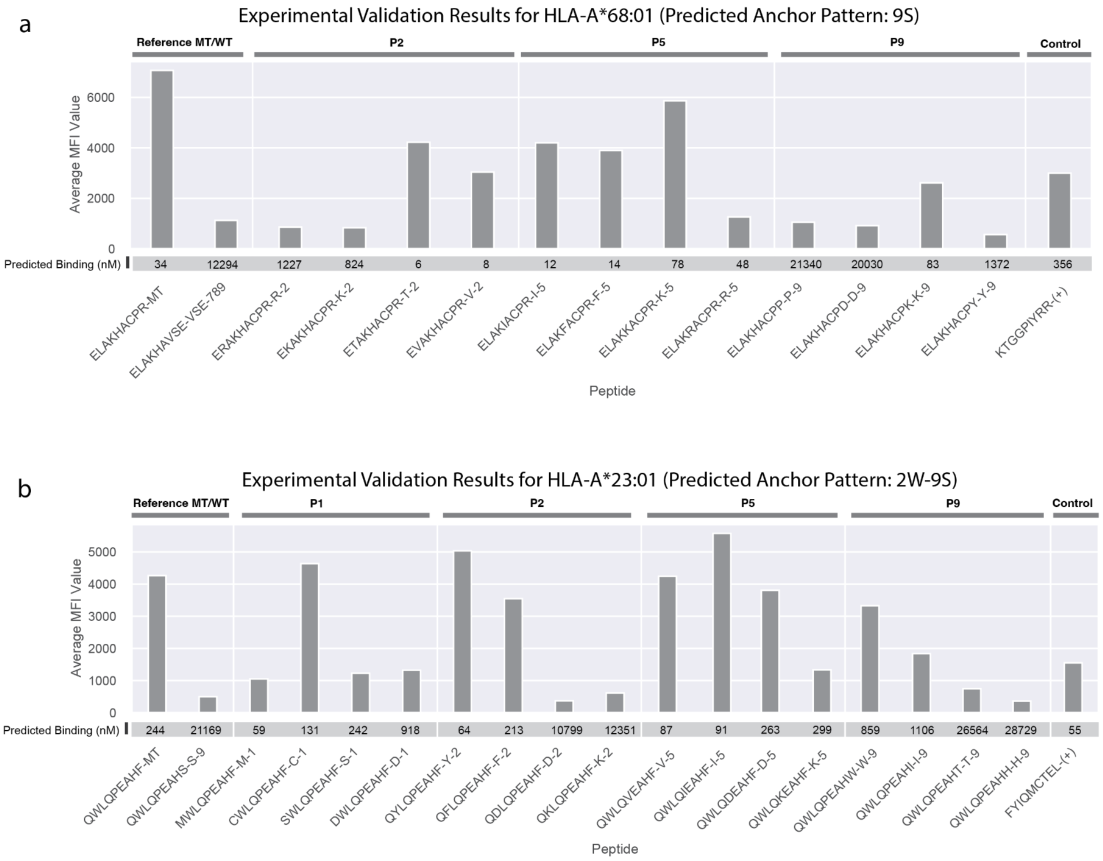

**Fig. S10. Breakdown of predicted binding affinity values versus measured binding affinity by individual algorithms**

Binding affinity values predicted by each individual algorithm plotted against the measured binding affinity (log[nM]). HLA-A*68:01, HLA-B*07:02, HLA-B*08:01, HLA-A*02:01 were subject to linear fitting with $R^2$ values shown (HLA-A*24:02 was excluded due to the limited range of data available). Non-binding peptide-HLA combinations were also excluded from this plot since they have no measured binding affinity value available. Prediction scores are shown for the following 8 different algorithms: MHCflurry, MHCnuggetsI, NetMHC, NetMHCcons, NetMHCpan, Pickpocket, SMM, and SMMPMBEC.

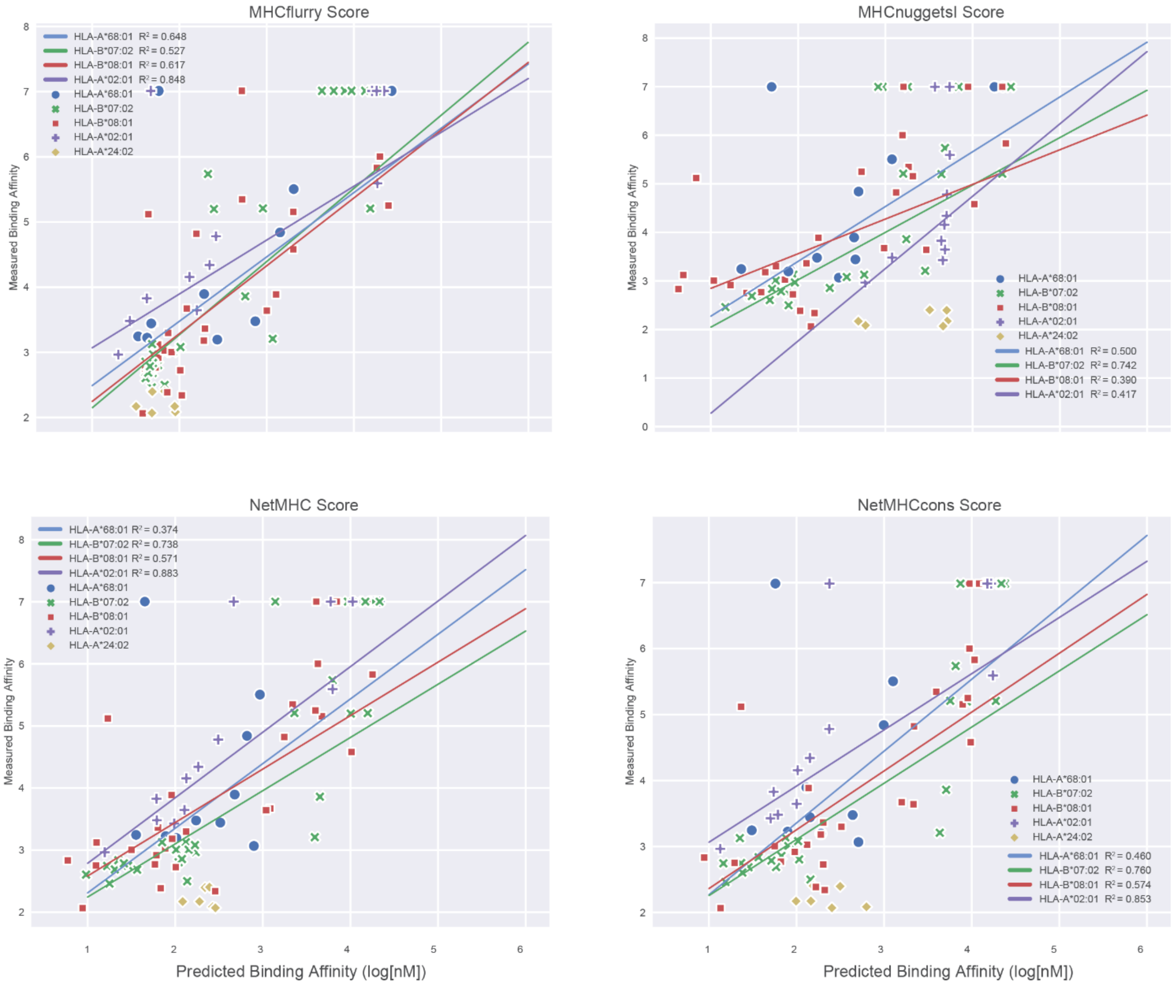

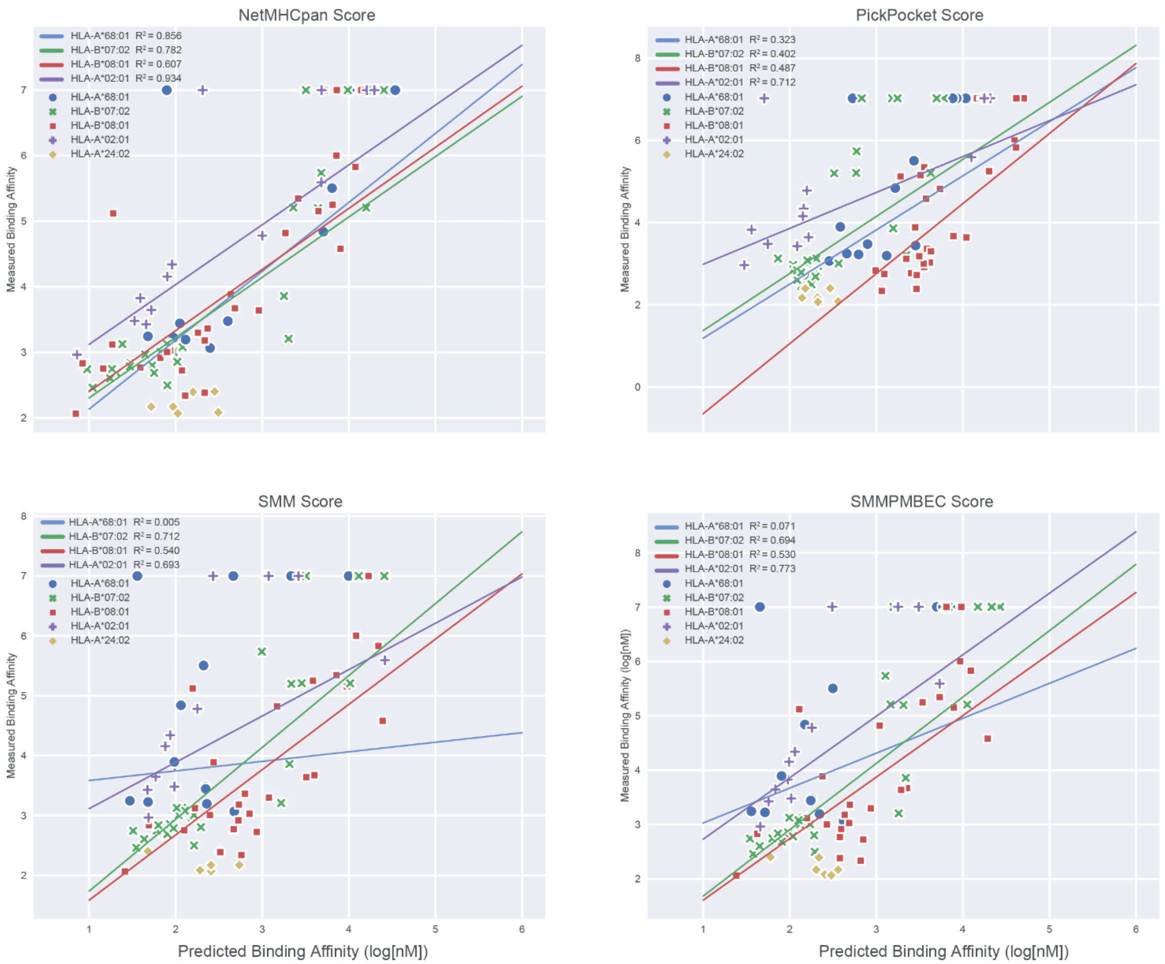

NetMHCpan Score
HLA-A*68:01 $R^2 = 0.856$
HLA-B*07:02 $R^2 = 0.782$
HLA-B*08:01 $R^2 = 0.607$
HLA-A*02:01 $R^2 = 0.934$
HLA-A*68:01
HLA-B*07:02
HLA-B*08:01
HLA-A*02:01
HLA-A*24:02
Measured Binding Affinity
PickPocket Score
HLA-A*68:01 $R^2 = 0.323$
HLA-B*07:02 $R^2 = 0.402$
HLA-B*08:01 $R^2 = 0.487$
HLA-A*02:01 $R^2 = 0.712$
HLA-A*68:01
HLA-B*07:02
HLA-B*08:01
HLA-A*02:01
HLA-A*24:02
Measured Binding Affinity
SMM Score
HLA-A*68:01 $R^2 = 0.005$
HLA-B*07:02 $R^2 = 0.712$
HLA-B*08:01 $R^2 = 0.540$
HLA-A*02:01 $R^2 = 0.693$
HLA-A*68:01
HLA-B*07:02
HLA-B*08:01
HLA-A*02:01
HLA-A*24:02
Measured Binding Affinity
Predicted Binding Affinity (log[nM])
SMMPMBEC Score
HLA-A*68:01 $R^2 = 0.071$
HLA-B*07:02 $R^2 = 0.694$
HLA-B*08:01 $R^2 = 0.530$
HLA-A*02:01 $R^2 = 0.773$
HLA-A*68:01
HLA-B*07:02
HLA-B*08:01
HLA-A*02:01
HLA-A*24:02
Measured Binding Affinity (log[nM])
Predicted Binding Affinity (log[nM])

**Fig. S11. Correlation scores between individual MHC binding algorithm predictions and measured binding affinities across HLA alleles**

Bar plot showing Pearson correlation scores calculated between predicted binding affinity values from 8 different algorithms and the measured IC50 binding affinity values for individual HLA alleles. Plots are grouped by the specific HLA allele and individual algorithms are marked with a color legend.

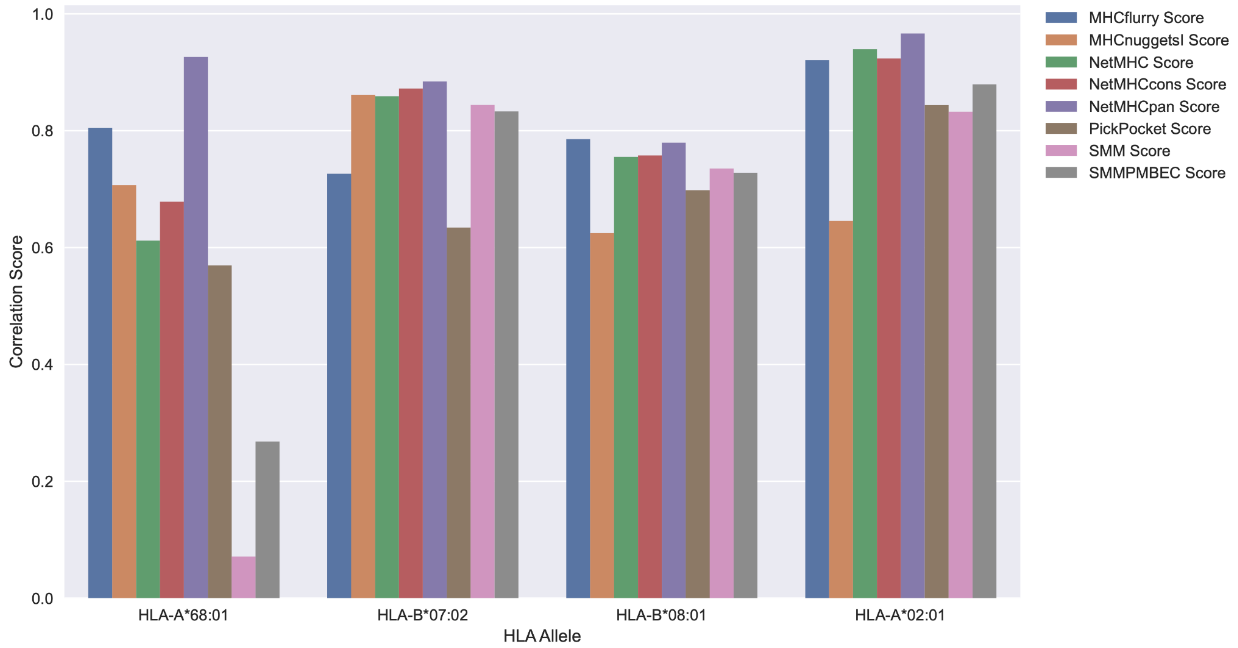

**Fig. S12. Distribution of anchor scenario categories for neoantigen candidates from 923 tumor-HLA paired samples**

Bar plot showing counts of neoantigen falling into each of the scenarios listed in Fig 7a. by analyzing neoantigen candidates from 923 tumor-HLA paired TCGA samples.

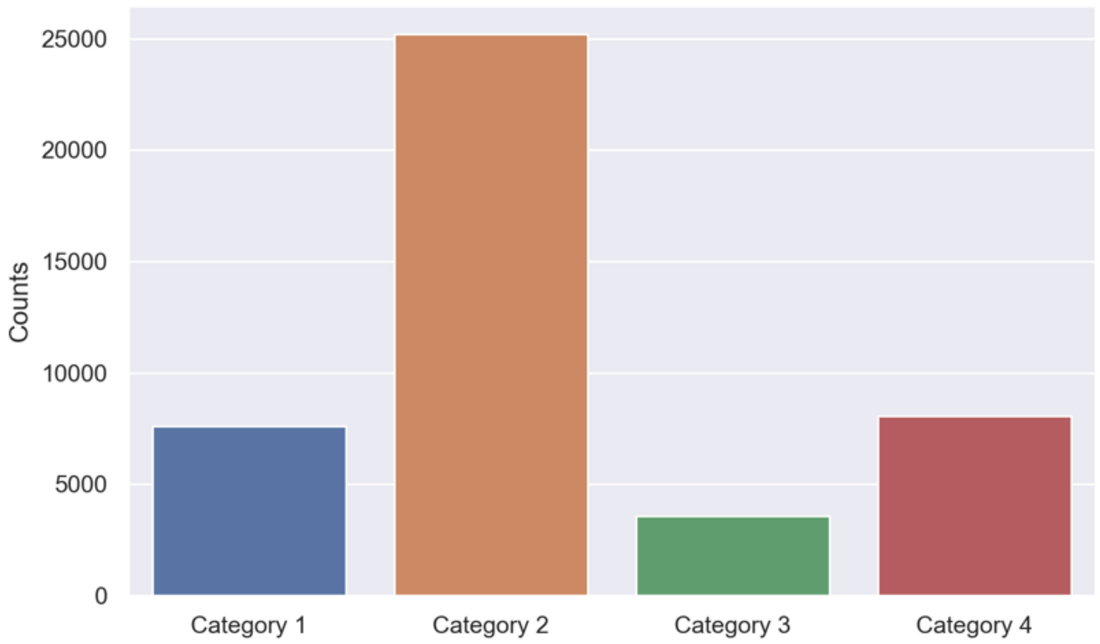

**Fig. S13. Patient-level analysis for impact of anchor considerations on neoantigen prioritization**

Patient-level impact of anchor position information on neoantigen prioritization decisions using 100 randomly selected samples from a pool of 1,356 TCGA samples. **a**, The distribution of tumor sample types for a TCGA sample pool (n = 1,356) is shown as a bar plot. The x-axis represents the different cancer types included and the y-axis shows the number of patient samples. **b**, A scatterplot shows the number of unique variants plotted against the number of predicted strong binding neoantigen candidates. Neoantigen candidates were compiled such that each variant had its top neoantigen candidate selected. The candidates were subsequently filtered based on a 500 nM binding affinity cutoff for each patient. The inner bar plot shows the distribution of tumor types for the randomly selected 100 patients. The legend shows the color labeling for each tumor type and is consistent between outer scatterplot and inner bar plot. **c, d, e:** Histogram plots showing distribution of neoantigen prioritization decision differences between filters A, B and C (Methods). Differences were normalized with respect to each individual patient's neoantigen counts and presented as the percentage of peptides that would be classified differently under each anchor interpretation scheme in terms of its neoantigen candidacy. The x-axis shows the range of percentage differences between filters while the y-axis shows patient sample counts for respective bins.

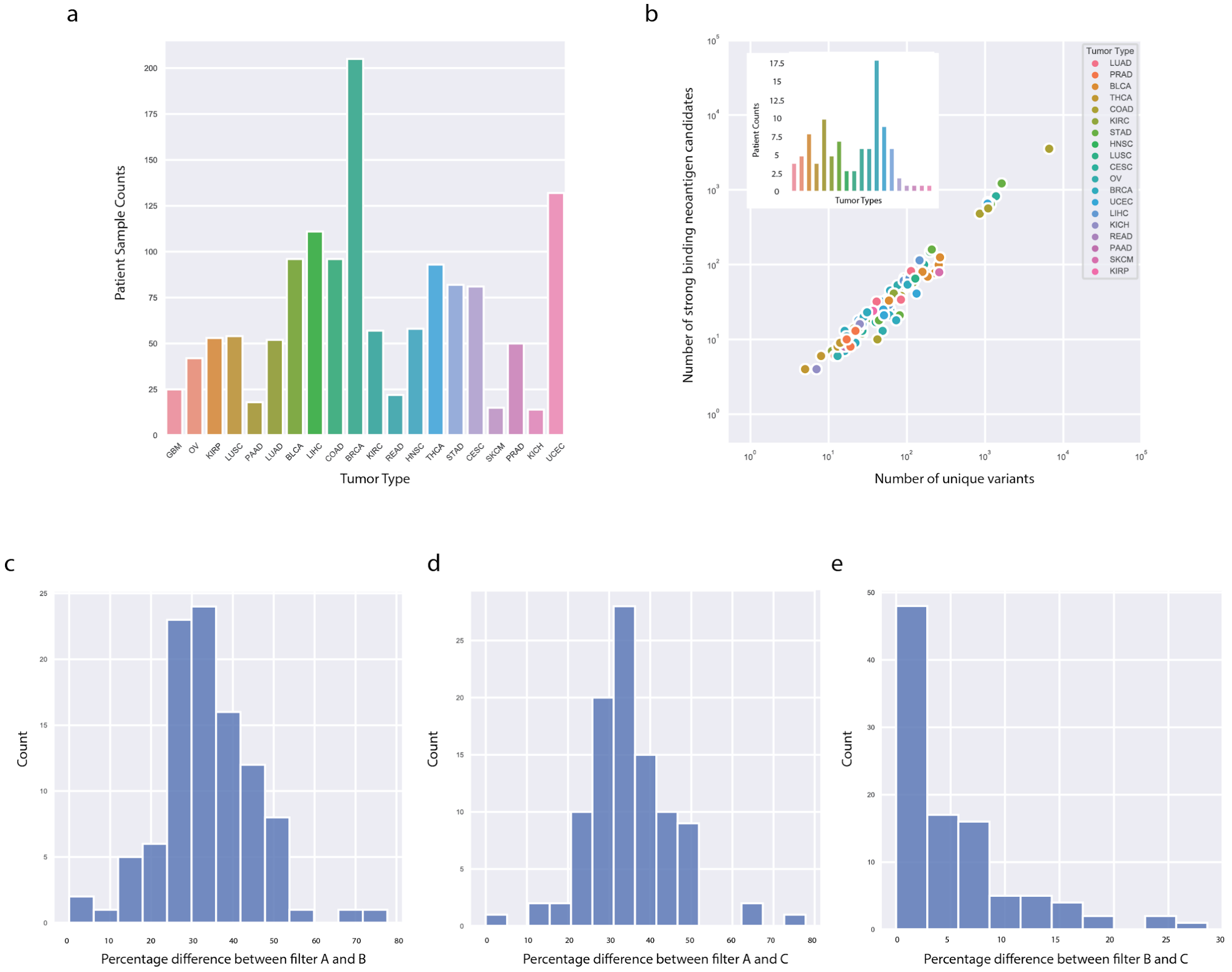

### Supplementary Tables

**Data file S1. Seed dataset of strong binding neoantigen candidates**
We identified 609,807 strong binding peptides for 328 HLA alleles from clinical and TCGA datasets. These served as a seed dataset for our anchor prediction workflow where a maximum of 10 peptides were selected at random for each HLA-peptide length combination. Peptide sequences, corresponding strong binding HLA alleles and the predicted median binding affinities across up to 8 different algorithms are included. The full dataset with all individual algorithmic prediction scores is also made available at http://genomedata.org/anchor_predictions/.

**Data file S2. Anchor predictions for 328 HLA alleles**
Prediction results from our computational workflow sorted by HLA alleles and peptide lengths. Overall scores are listed by peptide positions and represent the level of binding affinity change when mutated at the particular location. The 9-mer peptide data section contains additional information matching individual HLA alleles to their color-coded cluster in Figure 3.

**Data file S3. HLA Summary Information**
Information regarding all HLA alleles analyzed, including: 1) Nearest neighbor as defined by NetMHCpan, 2) Distance to nearest neighbor, 3) mean standard deviation of scores across all algorithms predicting for that allele using all available peptides in the seed dataset, 4) anchor cluster predicted as shown in Fig. 3., 5) training data available for all HLA alleles based on NetMHCpan4.0 (both elution and binding data) 6) number of algorithms able to generate predictions for each HLA allele, and 7) the detailed list of algorithms included in the previous column count. Note that if an HLA allele had training data, then it did not need to use a nearest neighbor for estimating binding and hence distance is 0 with nearest neighbor listed as itself.

**Data file S4. HLA PDB data table**
All protein data bank structures, collected for orthogonal validation of anchor prediction results, are listed. Table includes: the specific HLA allele and peptide pair, the PDB identifiers, predicted binding affinities and anchor cluster codes. The first tab of the spreadsheet contains information for structures collected that only contain the HLA-peptide complex and the second tab contains those collected that contain HLA-peptide-TCR complexes.

**Data file S5. Orthogonal validation correlation data**
A subset of X-ray crystallography structures were used in demonstrating the distribution of correlation scores for the distance and SASA metrics against prediction scores. Information on the subset data used such as peptide sequence, HLA allele, distance/SASA correlations and respective p values are included. The first tab of the spreadsheet contains information on the subset plotted from HLA-peptide only structures and the second tab contains information on the subset plotted from HLA-peptide-TCR complexes.

**Data file S6. Summary of all in vitro and cell based experimental validation data**
Validation experiments were performed on a total of 136 peptide-HLA combinations. These experiments include both IC50 binding assays (measured binding affinity) as well as cell-based stabilization assays (average MFI value). A summary of the entire dataset as well as information on predicted binding affinities and measured binding categories are included.

**Data file S7. Breakdown of individual algorithm predictions and their correlation with validation data**
Individual algorithm predictions (across 8 different algorithms) and their correlation with data from IC50 binding assays (measured binding affinity) were calculated. The spreadsheet includes two sheets, one with the individual algorithm scores for each peptide-HLA combination and the other with Pearson correlation coefficients for each algorithm across HLA alleles.

**Data file S8. List of TCGA samples for impact analysis**
A subset of TCGA samples were chosen using HLA-balance-based selection for our overall cohort-level impact analysis (Tab 1). An additional 100 TCGA samples were selected at random to further evaluate patient-level impact of anchor considerations (Tab 2). Specific TCGA sample names are included.

### Supplementary Video

**Movie S1. Demonstration of orthogonal validation using distance and SASA metrics from x-ray crystallography structures**
Video of the X-ray crystallography structure of HLA-B*08:01 and 9-mer peptide FLRGRAYGL. The video highlights the three components of the complex, including HLA (green), peptide (pink) and B2M (blue). It also provides a zoomed in view of the MHC binding groove, showcasing atoms surrounding the peptide in spheres and sticks for spatial perspective.
